# Comparative analysis of Alzheimer’s disease knock-in model brain transcriptomes implies changes to energy metabolism as a causative pathogenic stress

**DOI:** 10.1101/2021.02.16.431539

**Authors:** Karissa Barthelson, Morgan Newman, Michael Lardelli

## Abstract

Energy production is the most fundamentally important cellular activity supporting all other functions, particularly in highly active organs such as brains. Here, we summarise transcriptome analyses of young adult (pre-disease) brains from a collection of eleven early-onset familial Alzheimer’s disease (EOfAD)-like and non-EOfAD-like mutations in three zebrafish genes. The one cellular activity consistently predicted as affected by only the EOfAD-like mutations is oxidative phosphorylation that produces most of the brain’s energy. All the mutations were predicted to affect protein synthesis. We extended our analysis to knock-in mouse models of *APOE* alleles and found the same effect for the late onset Alzheimer’s disease risk allele ɛ4. Our results support a common molecular basis for initiation of the pathological processes leading to both early and late onset forms of Alzheimer’s disease and illustrate the utility of both zebrafish and knock-in, single EOfAD mutation models for understanding the causes of this disease.

## Introduction

Alzheimer’s disease (AD) is a complex and highly heterogenous neurodegenerative disease, defined the presence of intracellular neurofibrillary tangles (NFTs), and extracellular plaques consisting of a small peptide, amyloid β (Aβ) (Jack et al., 2018). AD was discovered over 100 years ago (Alzheimer, 1906). However, an effective therapeutic to halt, or even to slow disease progression, remains to be developed.

AD has a strong genetic basis (reviewed in (Sims et al., 2020)). In some rare cases, early-onset familial forms of AD (EOfAD, occurring before 65 years of age) arise due to dominant mutations in one of four genes: *PRESENILIN 1* (*PSEN1*), *PRESENILIN 2* (*PSEN2*), *AMYLOID β PRECURSOR PROTEIN* (*APP*), and *SORTILIN-RELATED RECEPTOR 1* (*SORL1*) (reviewed in (Barthelson et al., 2020a; Bertram and Tanzi, 2012; Temitope et al., 2021)). However, most AD cases are sporadic, showing symptom onset after the arbitrarily defined threshold of 65 years (late-onset sporadic AD, LOAD). Genetic variants at many loci have been associated with increased risk of LOAD (Kunkle et al., 2019; Lambert et al., 2013). The most potent variant is the ɛ4 allele of *APOLIPOPROTEIN E* (*APOE*) (Farrer et al., 1997) that has been described as “semi-dominant” (Genin et al., 2011).

An understanding of the early cellular stresses on the brain that eventually lead to AD is necessary to advance the development of preventative treatments. This is difficult to achieve through studying living humans, as access to young, pre-symptomatic brains is limited. Imaging studies have implicated structural and functional changes to the brain long before diagnosis of overt AD (Iturria-Medina et al., 2016; Quiroz et al., 2015). However, brain imaging cannot provide detailed molecular information on these changes. Transcriptome analysis is, currently, the strategy that can provide the highest resolution molecular description of cells and tissues. However, transcriptome analyses of ante-mortem brains carrying EOfAD mutations can only be performed using brain tissue from animal models.

Our group has exploited the zebrafish to generate a collection of knock-in models of EOfAD-like mutations in order to analyse their young brain transcriptomes (Barthelson et al., 2021; Barthelson et al., 2020b; Barthelson et al., 2020c; Barthelson et al., 2020e; Hin et al., 2020a; Hin et al., 2020b; Jiang et al., 2020; Newman et al., 2019). Our experimental philosophy has been to replicate, as closely as possible, the single heterozygous mutation state of EOfAD in humans, thereby avoiding possibly misleading assumptions regarding the molecular mechanism(s) underlying the disease. Our overall goal has been to compare a broad-range of EOfAD-like mutations in a number of EOfAD genes to define their shared pathological effects in young adult brains where the long progression to AD begins. To assist in this definition (by exclusion), we also created non-EOfAD-like mutations in the same genes as negative controls, i.e. frameshift mutations in the presenilin genes that do not cause EOfAD (reviewed in (Jayne et al., 2016), the “reading frame-preservation rule”). Since EOfAD and LOAD present as a similar diseases (reviewed in (Blennow et al., 2006; Masters et al., 2015)), despite their different genetic underpinnings, understanding the molecular effects of heterozygosity for EOfAD mutations may also give insight into changes occurring in LOAD.

Here, we summarise our findings of brain transcriptome analyses of EOfAD-like mutations in the zebrafish orthologues of genes implicated in EOfAD: *psen1, psen2* and *sorl1*. EOfAD mutations also exist in *APP*. However, zebrafish express two *APP* “co-orthologous” genes, *appa* and *appb*, complicating analysis of single, heterozygous mutations. Therefore, we re-analysed the best available publicly accessible brain transcriptomic data from a knock-in model of *APP* mutations: the *App*^*NL-G-F*^ mouse. Finally, we compared whether the brain transcriptome changes occurring due to single heterozygous EOfAD-like mutations in zebrafish are similar to the changes occurring due to the strongest genetic risk factor for LOAD: the ɛ4 allele of *APOE*, using publicly available brain transcriptome data from a humanised *APOE* targeted-replacement mouse model (APOE-TR) (Sullivan et al., 1997). We identify changes to energy metabolism as the earliest detectable cellular stress due to AD mutations, and demonstrate that knock-in zebrafish models are valuable tools to study the earliest molecular pathological events in this disease.

## Results

We first collated our findings from our zebrafish models of EOfAD-like mutations in *psen1* (Barthelson et al., 2021; Hin et al., 2020a; Hin et al., 2020b; Newman et al., 2019), *psen2* (Barthelson et al., 2020c) and *sorl1* (Barthelson et al., 2020b; Barthelson et al., 2020e). An advantage of using zebrafish for RNA-seq analyses is minimisation of genetic and environmental noise through breeding strategies such as that shown in **Figure 1A**. Large families of synchronous siblings can consist of heterozygous mutant and wild type genotypes allowing direct comparisons of the effects of each mutation. So far, we have performed six brain transcriptomic analyses based on various breeding strategies (summarised in **Table 1** and **Supplemental File 1**). The detailed analyses can be found in the publications cited above. However, the outcomes are summarised below and in **Figure 1**.

**Table 1:**
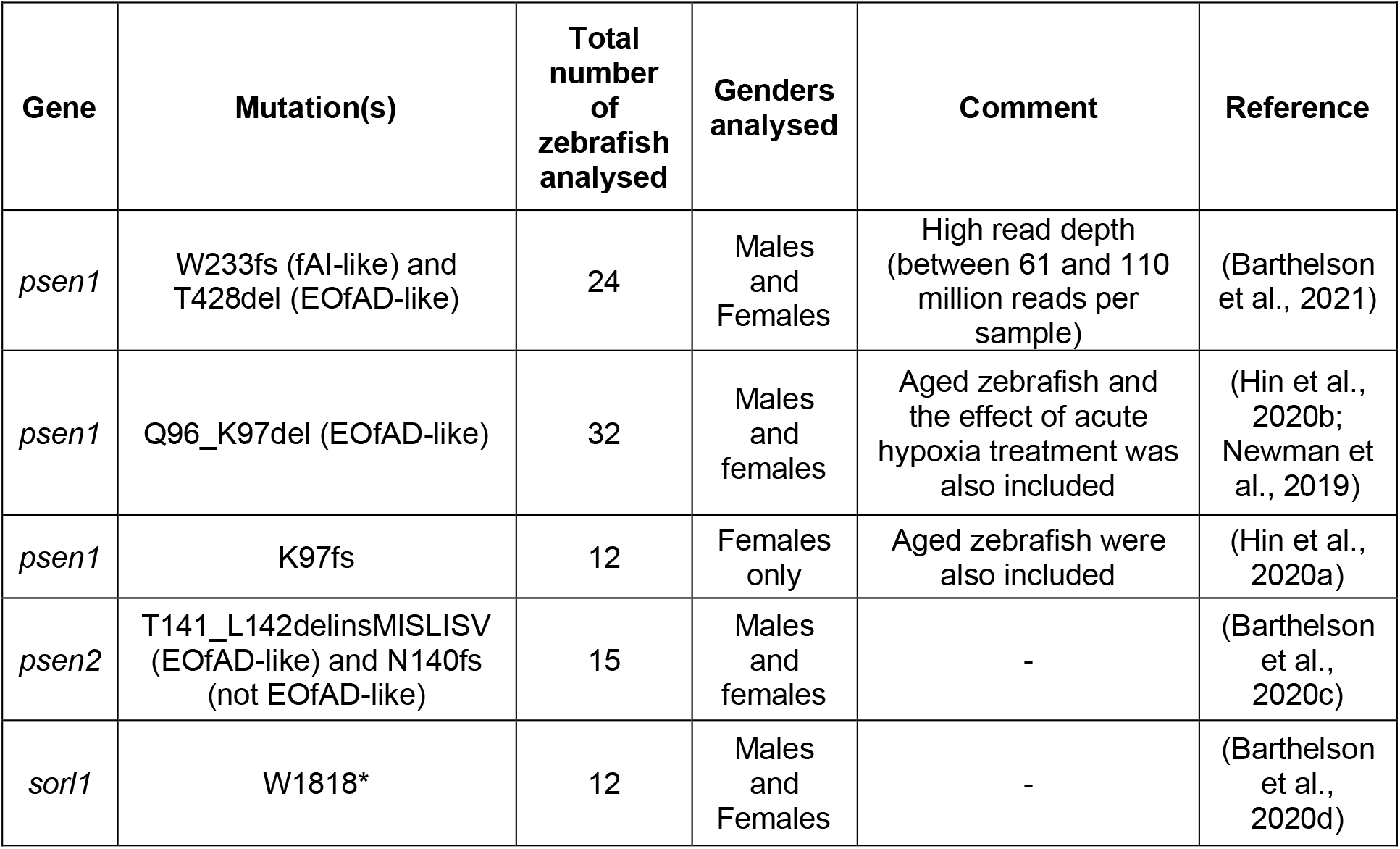
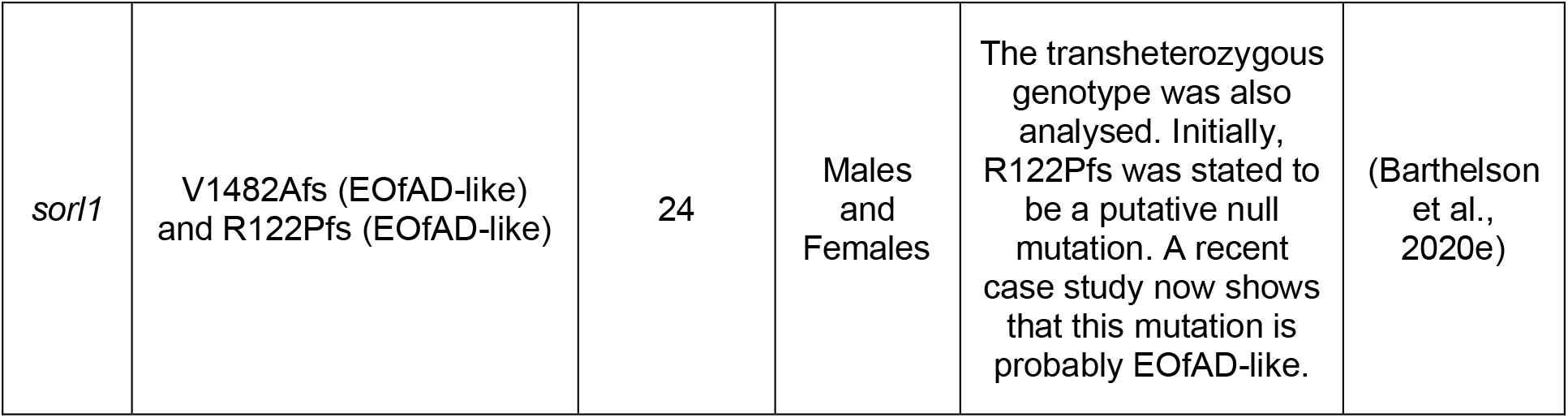
Summary of zebrafish RNA-seq experiments. For more detailed descriptions of study designs, see **Supplemental File 1**. EOfAD: early-onset familial Alzheimer’s disease. fAI: familial acne inversa.

**Figure 1:**
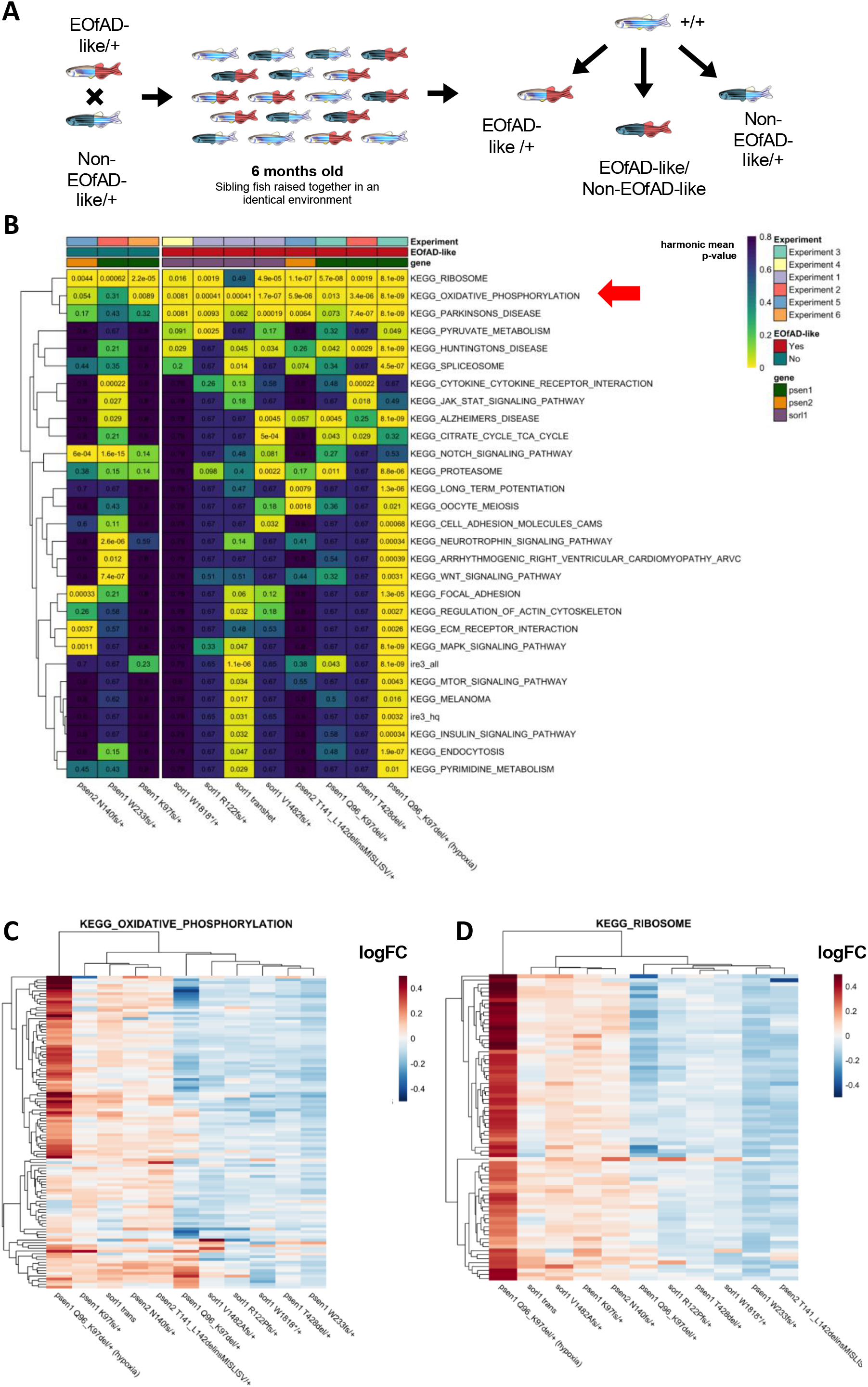
**A.** Schematic of a RNA-seq experiment using zebrafish. A single mating of a single pair of fish heterozygous for either an EOfAD-like or a non-EOfAD-like mutation results in a family heterozygous mutant, transheterozygous mutant, and wild type siblings. Comparisons made between genotypes in a RNA-seq experiment are depicted. **B.** Heatmap summary of significantly altered *KEGG* and *IRE* gene sets in zebrafish EOfAD genetic models at 6 months of age. Only gene sets significantly altered (FDR-adjusted harmonic mean p-value < 0.05) in at least two comparisons of mutant zebrafish to their corresponding wild type siblings are shown. Columns are grouped by whether or not the zebrafish genotype is EOfAD-like, while rows are clustered based on their Euclidean distance. The numbers are FDR-adjusted harmonic mean p-values. **C.** Heatmap indicating the logFC of genes in the *KEGG* gene sets for oxidative phosphorylation and, **D,** the ribosome in zebrafish mutants compared to their wild type siblings. Rows and columns are clustered based on their Euclidean distance. Only genes considered detectable in all RNA-seq experiments are depicted. See **Supplemental File 1** for more information on individual study designs.

We found that heterozygosity for most of our EOfAD-like mutations does not result in many differentially expressed (DE) genes in young adult brains (Barthelson et al., 2021; Barthelson et al., 2020b; Barthelson et al., 2020c; Barthelson et al., 2020e) (as would be expected for modelling a disease that becomes overt in middle age). Therefore, we performed gene set enrichment analyses to predict which cellular processes were affected by each of the mutations in each experiment. We used the *KEGG* (Kanehisa and Goto, 2000) gene sets to determine whether changes to gene expression were observed in any of 186 biological pathways/processes. Additionally, we recently proposed that neuronal iron dyshomeostasis may be an effect-in-common of EOfAD mutations in the context of AD pathogenesis (Lumsden et al., 2018). Therefore, we used our recently defined iron responsive element (*IRE*) gene sets (HIN ET AL., 2020B) to test for evidence of iron dyshomeostasis. Biological processes found to be affected in at least two different zebrafish mutants are shown in **Figure 1B**. The one gene set consistently altered by all of the EOfAD-like mutations, but not by the non-EOfAD-like mutations examined, is the *KEGG_OXIDATIVE_PHOSPHORYLATION* gene set, supporting that changes to mitochondrial function are an early cellular stress in EOfAD. The *KEGG_OXIDATIVE_PHOSPHORYLATION* gene set is also affected by heterozygosity for the K97fs mutation of *psen1*. K97fs is a frameshift mutation and so does not follow the “reading-frame preservation rule” (Jayne et al., 2016) of presenilin EOfAD mutations. However, K97fs (Hin et al., 2020a) models the PS2V isoform of human *PSEN2* (Sato et al., 1999), that shows increased expression in LOAD brains (see (Moussavi Nik et al., 2015) for an explanation) and so is still an AD-relevant mutation. Genes encoding the components of ribosomal subunits, as defined by the gene set *KEGG_RIBOSOME*, were affected by all the EOfAD-like mutations but also by non-EOfAD-like mutations in *psen1* and *psen2* (**Figure 1D**). Evidence for iron dyshomeostasis was also observed for the relatively severe EOfAD-like mutation *psen1*^*Q96_K97del*^/+ (under both normoxia and acute hypoxia conditions), and in transheterozygous *sorl1* mutants (i.e. with complete loss of wild type *sorl1*), as shown by significant changes to the expression of genes possessing IRE(s) in the 3’ untranslated regions of their encoded mRNAs.

EOfAD is also caused by mutations of the gene *APP.* Modelling of *APP* mutations in zebrafish is complicated by duplication of the *APP* orthologous gene in this organism. However, brain transcriptome data is available for a knock-in mouse model of EOfAD mutations in *APP*: the *APP*^*NL-G-F*^ mouse model (Castillo et al., 2017). In this model, the murine *APP* sequence is modified to carry humanised DNA sequences in the Aβ region, as well as the Swedish, Beyreuther/Iberian, and Arctic EOfAD mutations (Saito et al., 2014). While these mice do not closely reflect the genetic state of heterozygous human carriers of EOfAD mutations of *APP* (as the mice possess a total of six mutations within their modified *App* allele and are usually analysed as homozygotes), they should, at least, not generate artefactual patterns of gene expression change due to overexpression of transgenes (Saito et al., 2016). Castillo et al. (2017) performed brain transcriptomic profiling via microarray on the brain cortices of male homozygous *App*^*NL-G-F*^ mice relative to wild type mice at 12 months of age, as well as a transgenic mouse model of AD, 3xTg-AD mice (Oddo et al., 2003) relative to non-Tg mice. We re-analysed this microarray dataset to ask:

1. Are the *KEGG* gene sets affected in the male homozygous *App*^*NL-G-F*^ mice similar to those affected in EOfAD-like zebrafish?
2. Is there evidence for iron dyshomeostasis in the brains of these mice?

We did not observe alteration of any similar gene sets between the *App*^*NL-G-F*^ mice and our EOfAD-like zebrafish. However, any similarities may well have been masked by the overwhelming effects of greater age, variable environment (mouse litter-of-origin), and the effects of their six *App* mutations on brain cortex cell type proportions and inflammatory processes. The most statistically significantly affected cellular process in 12-month-old *App*^*NL-G-F*^ mice is lysosomal function as represented by the *KEGG_LYSOSOME* gene set (discussed later). Additionally, a plethora of inflammatory gene sets are also affected, with changes in the relative proportions of glial cells, particularly microglia, contributing to the appearance of increased levels of these gene transcripts in the bulk cortex RNA analysed (**Figure 2**).

**Figure 2:**
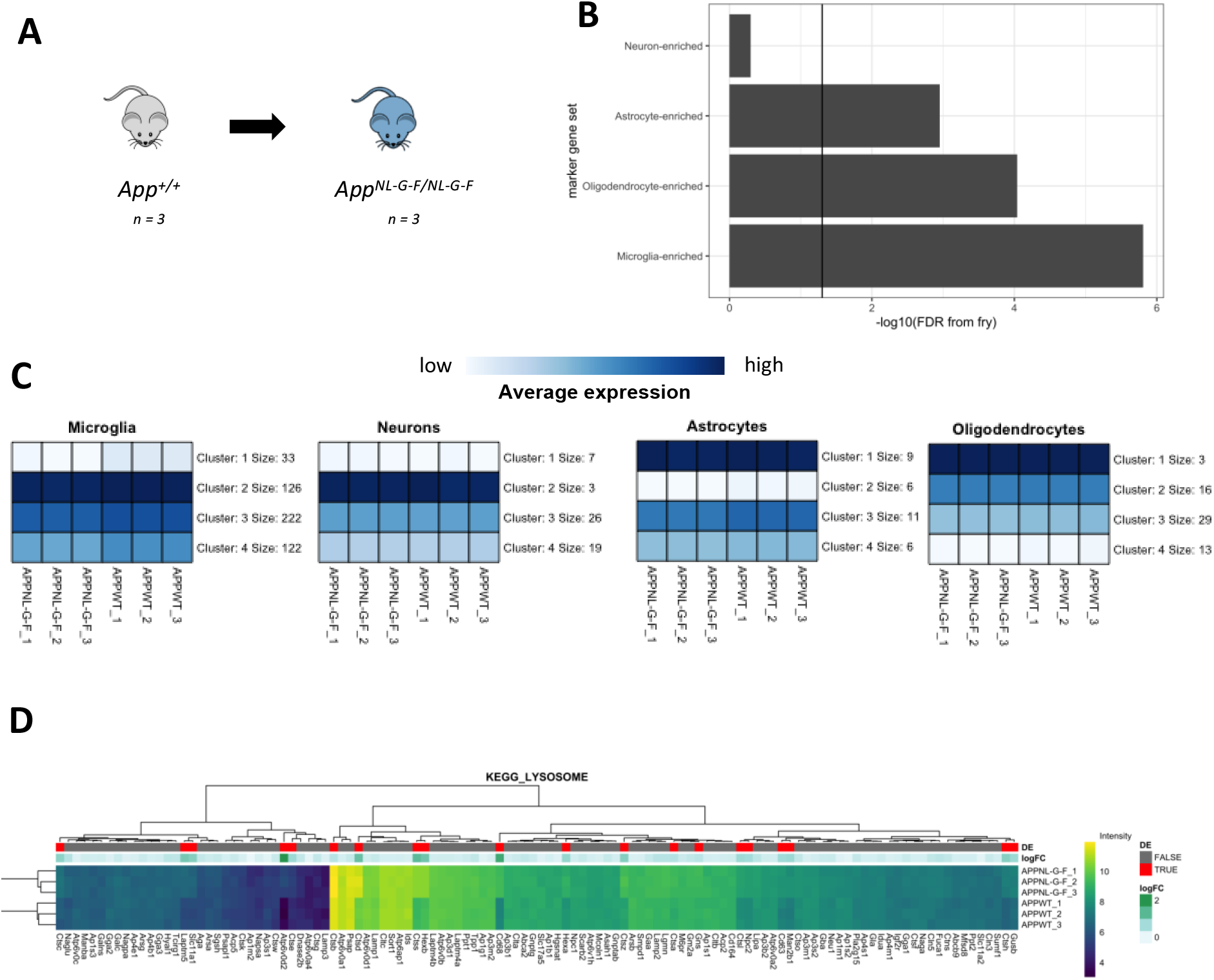
**A.** Visual representation of comparison of homozygous *App*^*NL-G-F*^ mice to wild type mice. **B.** Bar chart showing the FDR-adjusted p-value (directional hypothesis) from *fry* on marker genes of neurons, oligodendrocytes, astrocytes and microglia in *App*^*NL-G-F*^ relative to wild type. **C.** Heatmap indicating the expression of genes within these marker gene sets, summarised using K-means (K = 4). **D.** Heatmap showing the expression of genes in the *KEGG_LYSOSOME* gene set, clustered by their Euclidean distance. Each gene is labelled in red if it was identified as differentially expressed, and the magnitude of the fold change (logFC) is shown in green. See **Supplemental File 2** for more details.

A puzzling observation from the genome-wide association studies (GWAS) of LOAD, is that none of the risk variants identified fall within the EOfAD genes, *PSEN1*, *PSEN2*, or *APP* (Kunkle et al., 2019; Lambert et al., 2013). This has led to speculation that EOfAD and LOAD may be distinct diseases despite their histopathological and cognitive similarities (reviewed in (DeTure and Dickson, 2019; Tellechea et al., 2018)). Only one gene identified by GWAS of LOAD is suspected to harbour mutations causative of EOfAD, *SORL1*. Mutations in *SORL1* cause AD with ages of onset typically later than many mutations in *PSEN1* or *APP* (or may be incompletely penetrant) (Pottier et al., 2012; Thonberg et al., 2017). Nevertheless, as shown in **Figure 1**, we identified changes in the gene set *KEGG_OXIDATIVE_PHOSPHORYLATION* in young adult zebrafish heterozygous for EOfAD-like mutations in *sorl1,* as well as in zebrafish modelling overexpression of the PS2V isoform that is upregulated in LOAD. Therefore, we asked whether changes in oxidative phosphorylation might be associated with the most common genetic risk factor for LOAD, the ɛ4 allele of the gene *APOE* (Corder et al., 1993; Genin et al., 2011; Kunkle et al., 2019; Lambert et al., 2013; Saunders et al., 1993). Like *APP*, the *APOE* orthologous gene in zebrafish is refractory to analysis due to duplication. Therefore, to compare our zebrafish mutant data to early brain transcriptome changes caused by the ɛ4 allele of *APOE,* we analysed data from a set of human gene targeted replacement mouse models, APOE-TR (Sullivan et al., 1997). These mouse models transcribe human *APOE* alleles from the endogenous murine *Apoe* promotor: the predominant human allele ɛ3, the rare AD-protective ɛ2 allele, and the AD-risk allele, ɛ4. Zhao et al. (2020) performed a comprehensive brain transcriptome profiling experiment across aging in both male and female mice to assess the effect of homozygosity for the ɛ2 or ɛ4 alleles relative to the risk-neutral ɛ3 allele. Since our aim is to identify the early changes occurring due to AD-related mutations, we re-analysed only the 3-month brain samples from the Zhao et al. dataset (i.e. omitting the samples from 12- and 24-month-old mice). For details of this re-analysis see **Supplementary File 3**. We found statistical evidence for significant changes in expression of oxidative phosphorylation and ribosome gene sets in mice homozygous for the humanised ɛ4 *APOE* allele, consistent with our zebrafish models of EOfAD. These effects were not observed for the AD-protective ɛ2 allele (**Figure 3**). Note that the effects of *APOE* genotype were highly dependent on the litter-of-origin of the samples and that changes to cell type proportions were observed in the male APOE4 mice. Therefore, future replication of this analysis with better-controlled transcriptome data is desirable to confirm that the effects observed are due to the *APOE* genotype.

**Figure 3:**
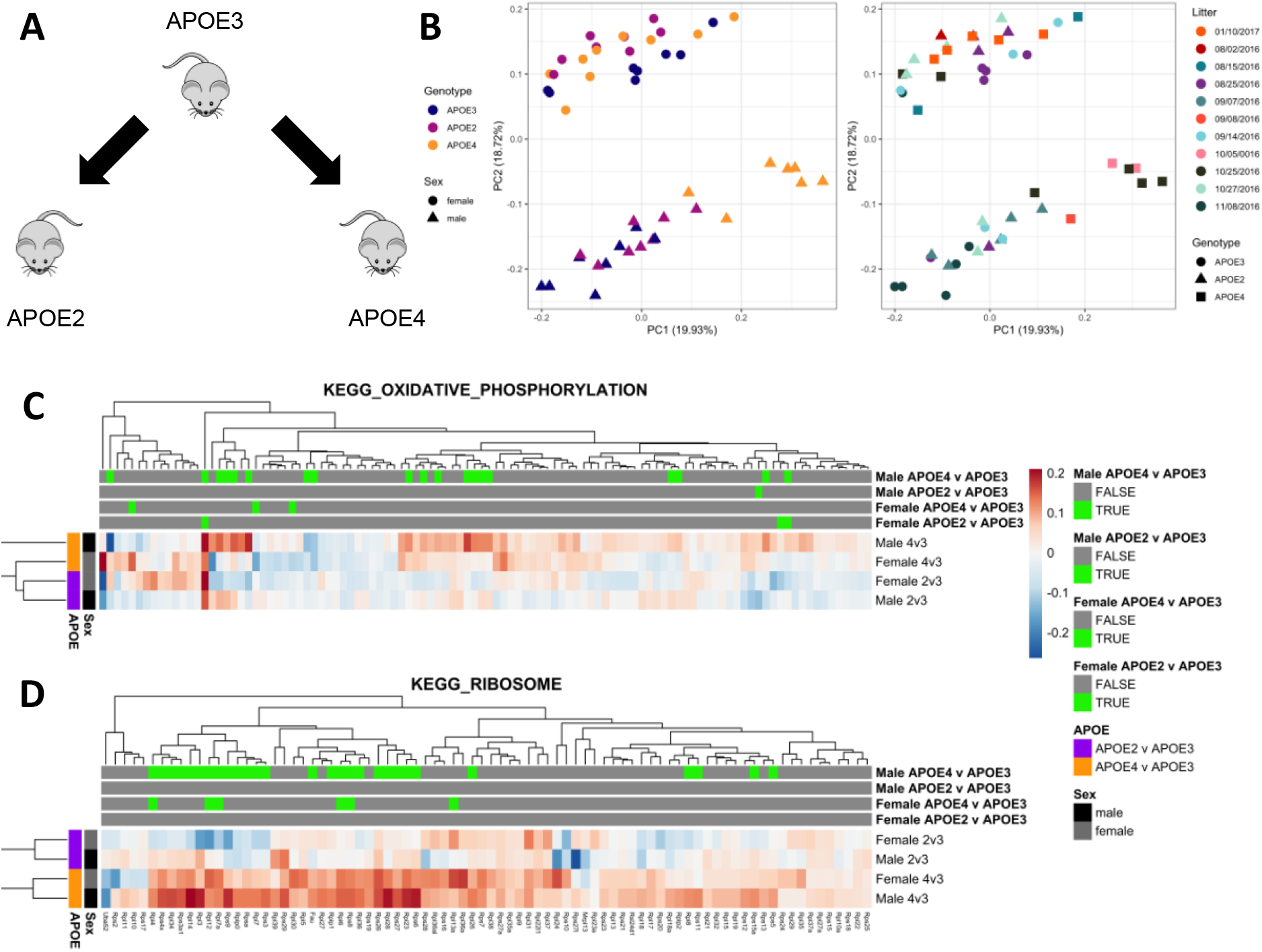
**A.** Visual representation of comparison of APOE4 or APOE2 mice to APOE3. This was performed for both male and female mice separately. **B.** Principal component analysis (PCA) of three month old APOE-TR mice. Principal component 1 (PC1) is plotted against PC2. The numbers between parentheses indicate the percentage of variation in the dataset explained by a PC. In the left graph, each point represents a sample, which is coloured by *APOE* genotype, and shaped by sex. In the right plot, each point is coloured according to litter (implied from the date of birth of each mouse), and shaped by *APOE* genotype. **C** Heatmap showing the logFC of genes in the *KEGG_OXIDATIVE_PHOSPHORYLATION* and **D** *KEGG_RIBOSOME* gene sets in APOE-TR mice. Genes are labelled in green above whether they were classified as differentially expressed (FDR < 0.05) in the differential gene expression analysis in a particular comparison. See **Supplemental File 3** for more details.

## Discussion

Energy production is the most fundamental of cellular activities. Life cannot be sustained without energy, and all other cellular activities depend upon it. The human brain, in particular, has very high energy demands and consumes the majority of the body’s glucose when at rest (reviewed in (Zierler, 1999)). Within the brain, the majority of energy use is to maintain the Na^+^-K^+^ membrane potential of neurons (Attwell and Laughlin, 2001) and neurons are assisted in meeting these energy demands by support from, primarily, astrocytes (e.g. via the astrocyte–neuron lactate shuttle (Pellerin and Magistretti, 1994)). All cells allocate considerable portions of their energy budgets to protein synthesis to maintain their structure and activity (Buttgereit and Brand, 1995). Energy is also required to maintain the low pH and high Ca^2+^ concentration of the lysosome (Christensen et al., 2002), the organelle which mediates uptake and recycling (autophagy) of cellular structural constituents (e.g. the amino acids for protein synthesis) (reviewed in (Yim and Mizushima, 2020)), Lysosomes are important for uptake and recycling of ferrous iron (Yambire et al., 2019), that is essential for oxidative phosphorylation by mitochondria (Oexle et al., 1999). On the lysosomal membrane, mTOR complexes sense nutrient and energy status to regulate protein synthesis, autophagy, and mitochondrial activity (reviewed in (Lim and Zoncu, 2016)).

Our analyses of young adult brain transcriptomes in zebrafish have found that five EOfAD-like mutations in a total of three EOfAD gene orthologues (*psen1*, *psen2*, and *sorl1*) all cause statistically significant effects on the expression of genes involved in oxidative phosphorylation, while non-EOfAD-like mutations in *psen1* and *psen2* do not. Therefore, effects on oxidative phosphorylation are a common, early “signature” of EOfAD. Intriguingly, we previously observed downregulation of the oxidative phosphorylation genes due to heterozygosity for the *psen1*^Q96_K97del^ mutation in whole zebrafish larvae at 7 days post fertilisation (dpf) (Dong et al., 2020), suggesting changes to mitochondrial function are a very early cellular stress in AD pathogenesis. All of the mutations studied have also resulted in changes in the levels of transcripts required for ribosome formation and we suggest this may reflect changes in mTOR activity (see **Supplementary File 4**). The reason for the different effects of EOfAD-like and non-EOfAD like mutations in these genes is uncertain. However, for the *PRESENILIN*s, the single most consistent characteristic of the hundreds of known EOfAD mutations is that they maintain the ability of the genes to produce at least one transcript isoform with the original reading frame (the “reading frame preservation rule” (Jayne et al., 2016)). This strongly supports that all these mutations act via a dominant gain-of-function molecular mechanism (to interfere with a normal cellular function). The EOfAD genes, *PSEN1*, *PSEN2*, *APP*, and *SORL1*, encode proteins which are all expressed within the endolysosomal pathway of cells (Andersen et al., 2005; Kawai et al., 1992; Pasternak et al., 2003; Sannerud et al., 2016) and also within the mitochondrial associated membranes (MAM) of the endoplasmic reticulum (Area-Gomez et al., 2009; Lim, 2015). MAM are responsible for regulation of ATP production (through Ca^2+^ signalling (Duchen, 1992)), oxidative protein folding (reviewed in (Simmen et al., 2010)), and the initiation of autophagy (Hamasaki et al., 2013). Interestingly, like mutations in *PSEN1* (Lee et al., 2010) and the C99 fragment of APP (Jiang et al., 2019), the ɛ4 allele of APOE has also been shown to affect lysosomal pH (Prasad and Rao, 2018) and the MAM (Tambini et al., 2016). From the work in this paper, we have now seen that the “semi-dominant” ɛ4 LOAD risk allele (Genin et al., 2011), like EOfAD mutations, also affects the expression of genes involved in oxidative phosphorylation and ribosome function in young adult brains. Thus, early effects on oxidative phosphorylation (and so energy production) appear to be a common, early disturbance associated with both early- and late- onset forms of Alzheimer’s disease. Our analysis supports that the two diseases are part of a single pathological continuum, with age of onset likely influenced by the severity of the energy effect involved. (After all, the 65 year age threshold defined as differentiating “early-onset” from “late-onset” is arbitrary.) The mystery of why GWAS has failed to detect variation in *PSEN1*, *PSEN2*, or *APP* in LOAD remains, although some mutations in *PSEN2* and *APP* genes do cause later onset familial forms of the disease (Cruchaga et al., 2012) and/or show incomplete penetrance (Finckh et al., 2000; Rossor et al., 1996; Sherrington et al., 1996; Thordardottir et al., 2018) or recessive inheritance (Di Fede et al., 2009; Tomiyama et al., 2008).

Knock-in mouse models of single EOfAD mutations were generated 15-20 years ago (Guo et al., 1999; Kawasumi et al., 2004) but their brain transcriptomes have never been analysed in detail. This is likely because these mice showed only very mild cognitive phenotypes and lacked the AD histopathology currently used to define the disease (Aβ deposition and neurofibrillary tangles of tau protein (Jack et al., 2018)). By expressing multiple mutant forms of EOfAD genes in transgenic mice, Aβ plaques can be detected and cognitive changes observed (reviewed in (Esquerda-Canals et al., 2017; Myers and McGonigle, 2019)). However, experience with use of many such “mouse models of Alzheimer’s disease” has shown a lack of correlation of cognitive changes with Aβ levels (Foley et al., 2015) and transcriptome analysis of their brains has shown little to no concordance with transcriptomes from post-mortem AD brains, or between the models themselves (Hargis and Blalock, 2017). In two papers in 2014 and 2016, Saito and colleagues described phenotypic disparities between transgenic and *APP* EOfAD mutation knock-in mouse models (Saito et al., 2014; Saito et al., 2016). In the 2016 paper they went so far as to declare that,

> “*We recently estimated using single App knock-in mice that accumulate amyloidβ peptide without transgene overexpression that 60% of the phenotypes observed in Alzheimer’s model mice overexpressing mutant amyloid precursor protein (APP) or APP and presenilin are artifacts (Saito et al., 2014). The current study further supports this estimate by invalidating key results from papers that were published in Cell. These findings suggest that more than 3000 publications based on APP and APP/PS overexpression must be reevaluated*.”

Nevertheless, since 2016 thousands more papers have been published using transgenic mouse models of Alzheimer’s disease. In this light, we were surprised to find that *App*^*NL-G-F*^ homozygous mice display a young adult brain transcriptome that is more severely disturbed than in the multiply transgenic 3xTg-AD model - although that apparent disturbance is likely somewhat artefactual and likely due to changes in the relative proportions of different cell types in the model. Changes to cell type proportions are not observed in our zebrafish models (Barthelson et al., 2021; Barthelson et al., 2020c; Barthelson et al., 2020d, e; Hin et al., 2020a; Hin et al., 2020b), revealing another advantage of exploiting zebrafish in transcriptome analysis of bulk RNA from brain tissue (their highly regenerative brains (Kroehne et al., 2011) are likely more resistant to changes in cell type proportions).

Frustration with the difficulties of exploiting both transgenic and knock-in models of EOfAD mutations in mice has contributed to the drive for examining knock-in mouse models of LOAD risk variants, such as now conducted by the MODEL-AD Consortium (Oblak et al., 2020). The brain transcriptome similarities seen between our single mutation, heterozygous EOfAD mutation-like knock-in zebrafish models and the knock-in APOE ɛ4 mice strongly support the informative value of these models and imply that heterozygous EOfAD mutation knock-in mouse models offer a path forward, particularly in understanding the earliest molecular events that lead to Alzheimer’s disease.

## Acknowledgements

The authors would like to thank Dr. Nhi Hin for providing the Q96_K97del zebrafish dataset and original analysis. The results for the young APOE-TR mice were based on data obtained from the AD Knowledge Portal (https://adknowledgeportal.synapse.org/). Support for these data was provided by the NIH RF1 AG051504 and P01 AG030128. We thank Drs. Patrick Sullivan and Nobuyo Maeda for generating human APOE targeted replacement mice and providing access through Taconic.

## Author Contributions

K.B. performed the analysis. M.L. and M.N. conceived the project. All authors contributed to drafting and editing the manuscript.

## STAR Methods

### RESOURCE AVAILABILITY

#### Lead Contact

Further information and requests for resources and reagents should be directed to and will be fulfilled by the Lead Contact, Karissa Barthelson.

#### Materials Availability

This study did not generate new unique reagents.

#### Data and Code Availability Statement

The complete list of datasets (with links), software, and relevant algorithms used in this study is provided in the Key Resources Table. The code used to perform the analysis in this study can be found at https://github.com/karissa-b/AD-signature

### METHOD DETAILS

#### Zebrafish analysis

The harmonic mean p-values and differential gene expression analysis outputs were obtained from each individual analysis (see the Key Resources Table and (Barthelson et al., 2021; Barthelson et al., 2020b; Barthelson et al., 2020c; Barthelson et al., 2020e)). For these analyses, differential gene expression analysis was performed using *edgeR* (Robinson et al., 2009), and enrichment analysis was performed by calculation of the harmonic mean p-value (Wilson, 2019) of the raw p-values of three methods of ranked-list based enrichment analyses: *fry* (Wu et al., 2010), *camera* (Wu and Smyth, 2012), and *GSEA* (Subramanian et al., 2005) (as implemented in the *fgsea* R package (Sergushichev, 2016)). A gene set to be significantly altered if the FDR-adjusted harmonic mean p-value remained below 0.05 after FDR adjustment. The gene sets used for enrichment analysis were the *KEGG* (Kanehisa and Goto, 2000) gene sets, to determine whether changes to gene expression were observed in any of 186 biological pathways/processes. Additionally, we used our recently defined iron responsive element (*IRE*) gene sets (HIN ET AL., 2020B) to test for evidence of iron dyshomeostasis. For the K97fs and Q96_K97del analyses, enrichment analysis was not performed on the KEGG gene sets in the original analyses. Therefore, we performed the enrichment analysis as described above for these datasets. For the K97fs analysis, we obtained the gene-level counts and the results of the differential gene expression analysis described in (Hin et al., 2020a) from https://github.com/UofABioinformaticsHub/k97fsZebrafishAnalysis. For the Q96_K97del analysis, we obtained the gene-level counts and the and the results of the differential gene expression analysis described in (Hin et al., 2020b) from the first author of the cited paper.

#### *App*^*NL-G-F*^ microarray re-analysis

The detailed analysis can be found in **Supplemental File 2**, while the R code to reproduce the analysis can be found https://github.com/karissa-b/AD-signature. Briefly, the raw .CEL files (obtained from GEO) were imported into analysis with *R,* and pre-processed using the *rma* (Irizarry et al., 2003) method as implemented in the *oligo* package (Carvalho and Irizarry, 2010). We omitted any probesets which contained a median log2 intensity value of < 3.5 (lowly expressed) and also any probesets assigned to multiple genes. Differential gene expression analysis was performed using *limma* (Ritchie et al., 2015), specifying pairwise contrasts between the *App*^*NL-G-F*^ homozygous mice, or the 3xTg-AD homozygous mice with their respective controls by using a contrasts matrix. We considered a probeset to be differentially expressed in each contrast if the FDR adjusted p-value was < 0.05. For over-representation of the *KEGG* and *IRE* gene sets within the DE genes, we used *kegga* (Young et al., 2010). We also performed ranked-list based enrichment analysis as described for the zebrafish analysis.

#### APOE-TR RNA-seq re-analysis

The detailed analysis can be found in **Supplemental File 3**. We obtained the raw fastq files for the entire APOE-TR RNA-seq experiment from AD Knowledge Portal (accession number syn20808171, https://adknowledgeportal.synapse.org/). The raw reads were first processed using *AdapterRemoval* (version 2.2.1) (Schubert et al., 2016), setting the following options: *--trimns*, *--trimqualities, -- minquality 30, --minlength 35.* Then, the trimmed reads were aligned to the *Mus musculus* genome (Ensembl GRCm38, release 98) using *STAR* (version 2.7.0) (Dobin et al., 2013) using default parameters to generate .bam files. These bam files were then sorted and indexed using *samtools* (v1.10) (Li et al., 2009). The gene expression counts matrix was generated from the .bam files using *featureCounts* (version 1.5.2) (Liao et al., 2014). We only counted the number of reads which uniquely aligned to, strictly, exons with a mapping quality of at least 10 to predict expression levels of genes in each sample.

We then imported the output from *featureCounts* (Liao et al., 2014) for analysis with *R* (R Core Team, 2019). We first omitted genes which are lowly expressed (and are uninformative for differential expression analysis). We considered a gene to be lowly expressed if it contained, at most, 2 counts per million (CPM) in 8 or more of the 24 samples we analysed.

To determine which genes were dysregulated in APOE4 and APOE2 mice relative to APOE3, we performed a differential gene expression analysis using a generalised linear model and likelihood ratio tests using *edgeR* (McCarthy et al., 2012; Robinson et al., 2009). We chose a design matrix which specifies the *APOE* genotype and sex of each sample. The contrasts matrix was specified to compare the effect of APOE2 or APOE4 relative to APOE3 in males and in females. In this analysis, we considered a gene to be differentially expressed (DE) if the FDR-adjusted p-value was < 0.05. A bias for gene length and %GC content was observed in this dataset. Therefore, we corrected for this bias using conditional quantile normalisation (*cqn*) (Hansen et al., 2012). We calculated the average transcript length per gene, and a weighted (by transcript length) average %GC content per gene as input to *cqn* to produce the offset to correct for the bias. This offset was then included in an additional generalised linear model and likelihood ratio tests in *edgeR* with the same design and contrast matrices. For over-representation of the *KEGG* and *IRE* gene sets within the DE genes, we used *goseq* (Young et al., 2010), specifying average transcript length to generate the probability weighting function, which corrects for the probability that a gene is classified as DE based on its length alone.

We also performed ranked-list based enrichment analysis as described for the zebrafish analysis. Code to reproduce the analysis for the APOE-TR dataset can be found at https://github.com/karissa-b/AD-signature.

### QUANTIFICATION AND STATISTICAL ANALYSIS

#### Statistical analysis of *App*^*NL-G-F*^ microarray data

The detailed analysis can be found in **Supplemental File 2**. The sample numbers for the mouse *App*^*NL-G-F*^ microarray re-analysis consisted of n = 3 male mouse cortex transcriptomes per genotype (Castillo et al., 2017). The raw .CEL files were obtained from GEO, then imported into R and pre-processed using the *rma* method (Irizarry et al., 2003), which performs background correction, quantile normalisation and summarisation (median-polish). Differential gene expression analysis was conducted using *limma* v3.44.3 (Ritchie et al., 2015). The generated p-values were corrected for multiple testing using the false discovery rate (FDR). We considered a gene (probeset) to be differentially expressed if the FDR-adjusted p-value was less than 0.05 (no fold change cut-off).

Enrichment of the *KEGG* and *IRE* gene sets within the DE gene lists was performed using the *kegga.* We also performed ranked-list based enrichment analysis by calculation of the harmonic mean p-value (Wilson, 2019) of the raw p-values calculated from *fry* (Wu et al., 2010), *camera* (Wu and Smyth, 2012) and *fgsea* (Sergushichev, 2016). For enrichment analysis, we considered a gene set to be significantly altered if the FDR-adjusted p-value from *goseq*, or the FDR-adjusted harmonic mean p-value was less than 0.05.

#### Statistical analysis of APOE-TR RNA-seq data

The detailed analysis can be found in **Supplemental File 3**. Sample numbers mostly consisted of n = 8 mice per genotype, age and sex. However, upon inspection of the expression of genes of the Y-chromosome, we noted that three mice at 3 months of age had been incorrectly identified. The raw gene counts were calculated from *featureCounts,* normalised using the trimmed mean of M-values (TMM) method (Robinson and Oshlack, 2010), and used for calculation of gene expression values as logCPM (log2-counts-per-million). Differential gene expression analysis was performed using a generalised linear model and likelihood ratio tests as implemented in *edgeR* (Robinson et al., 2009). The generated p-values were corrected for multiple testing using the false discovery rate (FDR). We considered a gene to be differentially expressed if the FDR-adjusted p-value was less than 0.05 (no fold change cut-off). To test for over-representation of the *KEGG* and *IRE* gene sets within the DE genes, we used *goseq* (Young et al., 2010), specifying the average transcript length to calculate the probability weighting function. We considered a gene set to be significantly over-represented if the FDR-adjusted p-value generated by *goseq* was < 0.05. The ranked-list enrichment analysis was performed as described for the microarray data.

## Supplemental Information

**Supplemental File 1:** Zebrafish RNA-seq study designs

**Supplemental File 2:** App^NL-G-F^ microarray re-analysis. Related to Figure 2.

**Supplemental File 3:** APOE-TR RNA-seq re-analysis. Related to Figure 3.

**Supplemental File 4:** Additional discussion

