## Supplemental File 1 for "Comparative analysis of Alzheimer’s disease knock-in model brain transcriptomes implies changes to energy metabolism as a causative pathogenic stress"

***psen1* Q96_K97del experiment**

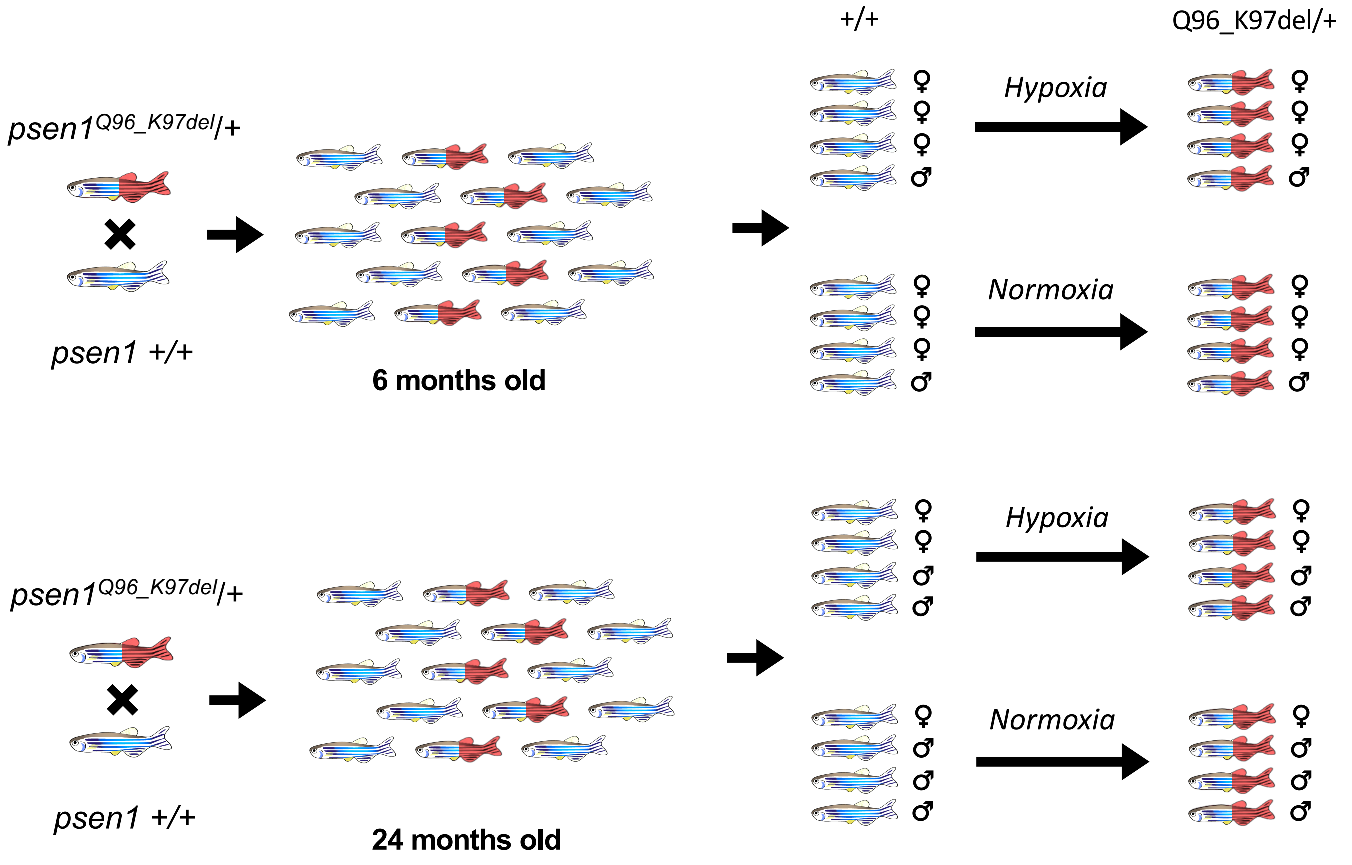


Two families of zebrafish were generated by mating a wild type fish with a fish heterozygous for the Q96_K97del mutation of *psen1*, resulting in families of fish either heterozygous for the Q96_K97del mutation, or wild type. These families were raised together in single tanks until 6 or 24 months of age. Then, subsets of the families were genotyped using allele-specific polymerase chain reactions (PCRs), followed by hypoxia treatment. Then fish were sacrified and n = 4 fish per genotype and treatment were subject to RNA-seq analysis.

***psen1* K97Gfs experiment**

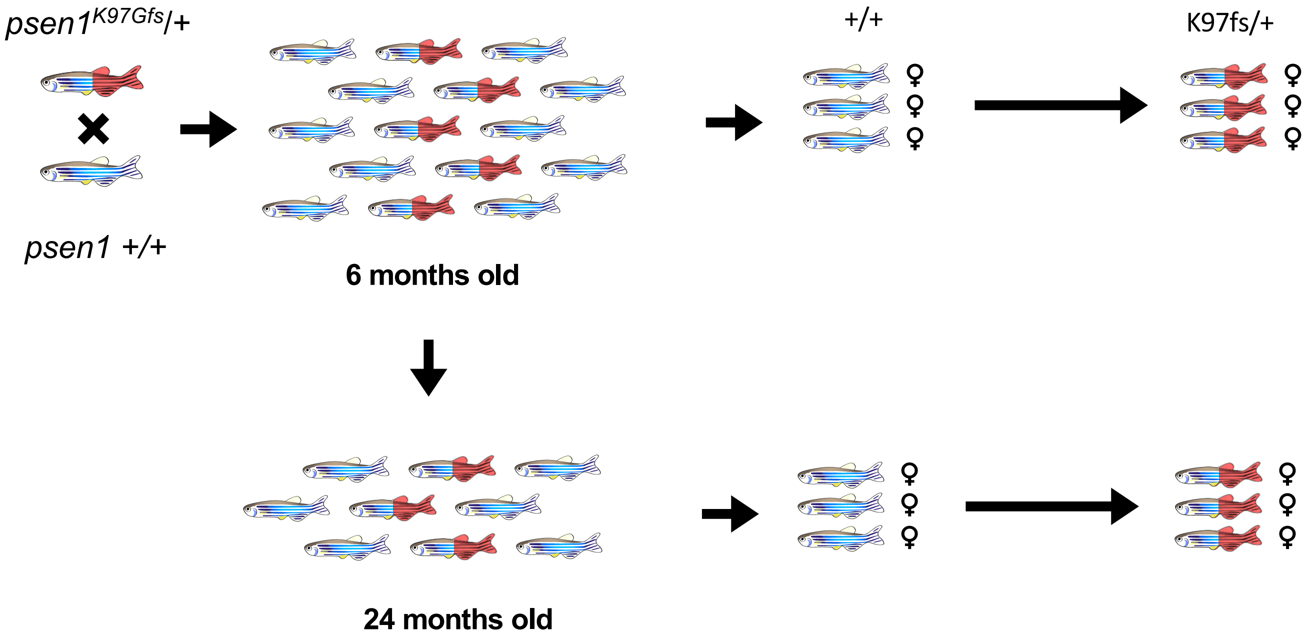


A wild type fish was mated with a fish heterozygous for the K97Gfs mutation of *psen1*, resulting in a family fish either heterozygous for the K97Gfs mutation, or wild type. This family of fish was raised together in a single tank until 6 months of age. Then, a subset of the family were genotyped using allele-specific polymerase chain reactions (PCRs), then fish were sacrificed and brains were removed for RNA-seq. The remaining fish in the tank were allowed to develop until 24 months of age where this was repeated to generate the aged samples for RNA-seq.

### *psen1* T428del vs W233fs experiment

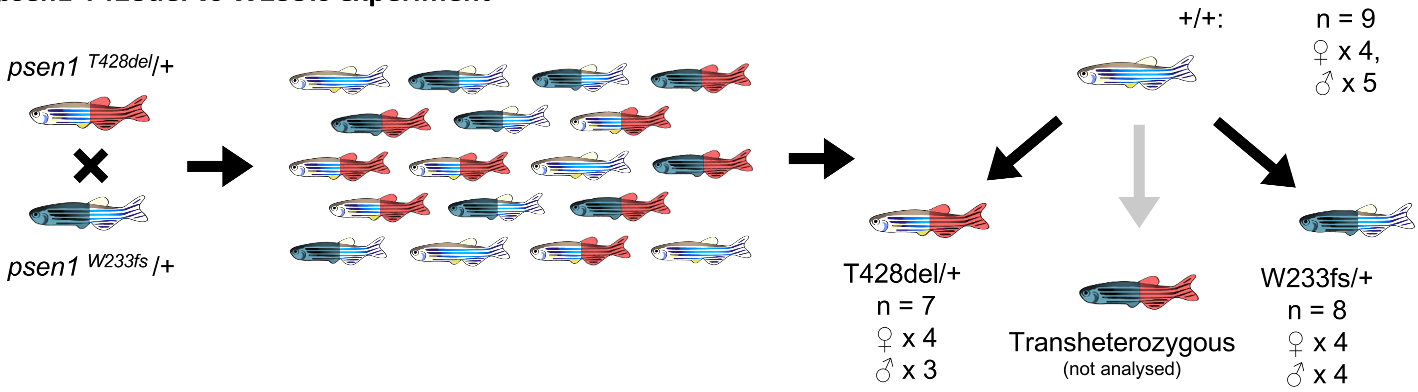


A fish heterozygous for the T428del (EOfAD-like) mutation of *psen1* was mated with a fish heterozygous for the W233fs mutation (similar to the P242fs mutation of human *PSEN1* causative for familial acne inversa) to generate a family of sibling fish with four possible *psen1* genotypes. This family was raised together in the same tank until 6 months of age, at which time 50 fish were randomly selected and sacrificed in a loose ice slurry. Fish were genotyped after sacrifice by allele specific PCRs. Then n = 8 fish per genotype (4 females and 4 males) were subjected to RNA-seq analysis. During the RNA-seq analysis, one T428del/+ fish was identified to be incorrectly genotyped and was re-classified as wild type.

### *psen2* frameshift vs EOfAD-like experiment

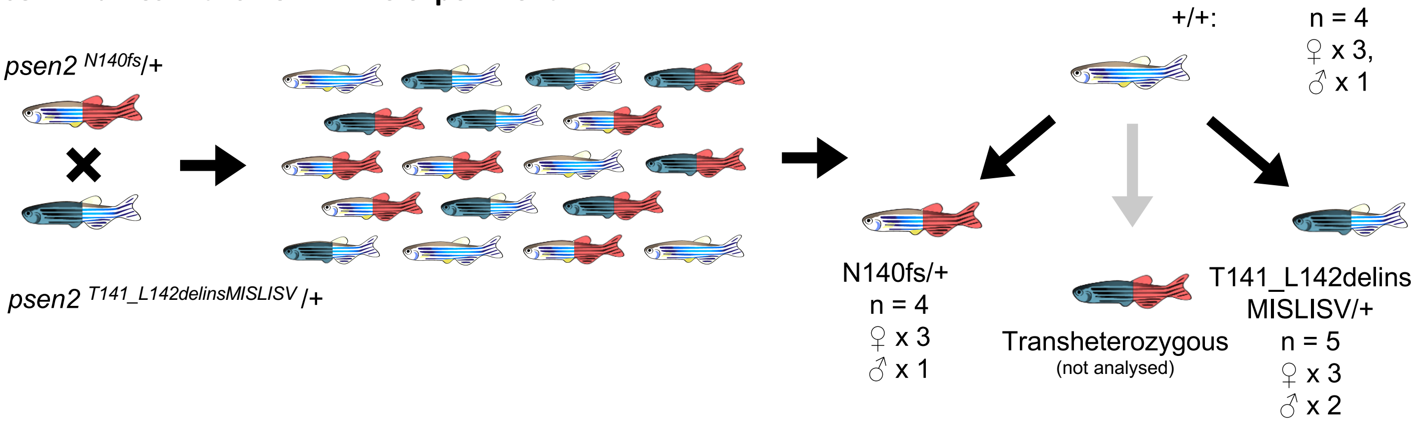


A fish heterozygous for the N140fs (not EOfAD-like) mutation of *psen2* was mated with a fish heterozygous for the T141_L142delinsMISLISV (EOfAD-like) mutation of *psen2* to generate a family of sibling fish with four possible *psen2* genotypes. This family was raised together in the same tank until 6 months of age, at which time 24 fish were randomly selected and sacrificed in a loose ice slurry (to allow for n = 5 of each genotype in the RNA-seq analysis). Fish were genotyped after sacrifice by allele specific PCRs. Then n = 5 fish per genotype (3 females and 2 males) were subjected to RNA-seq analysis. During the RNA-seq analysis, one wild type fish was an obvious outlier and was omitted from the rest of the analysis, and one N140s/+ sample has been incorrectly genotyped and was also omitted.

### *sorl1* W1818* experiment

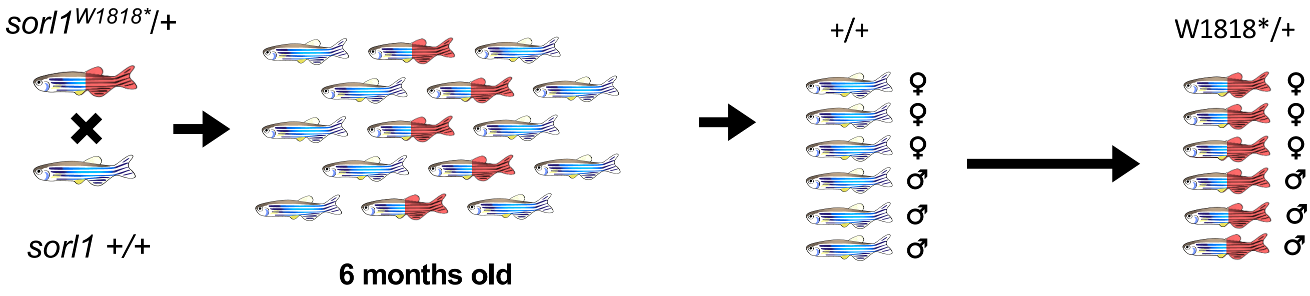


A fish heterozygous for the W1818* (EOfAD-like) mutation of *sorl1* was mated with a wild type fish to generate a family of sibling fish with two possible *sorl1* genotypes. This family was raised together in the same tank until 6 months of age, at which time 20 fish were randomly selected and sacrificed in a loose ice slurry (to allow for n = 6 of each genotype in the RNA-seq analysis). Fish were genotyped after sacrifice by allele specific PCRs. Then n = 3 fish per genotype and sex were subjected to RNA-seq analysis.

### *sorl1* V1482Afs and R122Pfs experiment

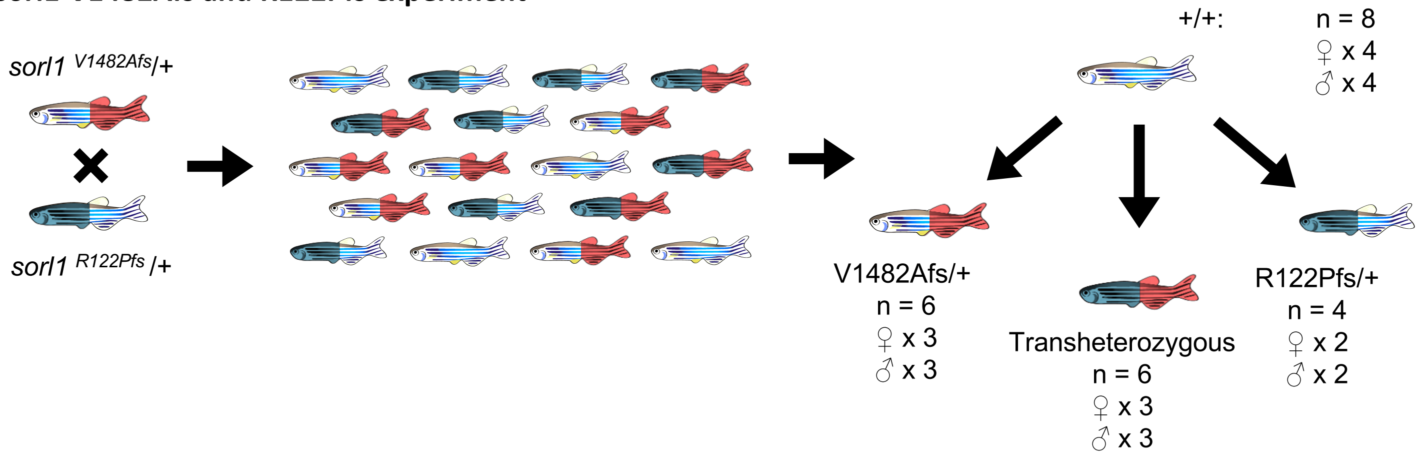


A fish heterozygous for the V1482Afs (EOfAD-like) mutation of *sorl1* was mated with a fish heterozygous for the R122Pfs mutation to generate a family of sibling fish with four possible *sorl1* genotypes. This family was raised together in the same tank until 6 months of age, at which time 50 fish were randomly selected and sacrificed in a loose ice slurry (to allow for n = 6 of each genotype in the RNA-seq analysis). Fish were genotyped after sacrifice by allele specific PCRs. During the RNA-seq analysis, two R122Pfs/+ fish were identified to be incorrectly genotyped and were re-classified as wild type.
