## Supplemental File 2 for "Comparative analysis of Alzheimer’s disease knock-in model brain transcriptomes implies changes to energy metabolism as a causative pathogenic stress"

Castillo et al. (2017) previously performed transcriptome analysis on the brains of two mouse models of AD: a knock-in model, *App*$^{NL-G-F}$ (Saito et al., 2014), and the transgenic model 3xTg-AD-H (Oddo et al., 2003). The *App*$^{NL-G-F}$ strain of mice carries a total of six mutations in the murine *App* gene: three mutations that humanise the mouse Aβ sequence, plus the Swedish (KM670/671NL), Iberian (I716F) and Artic (E693G) EOfAD mutations (Saito et al., 2014). The 3xTg-AD-H model of AD (hereafter referred to as 3xTg) carries the M146V EOfAD mutation within the endogenous mouse *Psen1* gene, and expresses two transgenes: human *APP* with the Swedish EOfAD mutation, and human *MAPT* with the P301L mutation. Transcription of the transgenes is driven by the mouse *Thy1.2* promotor (Oddo et al., 2003).

The cortices of three male mice of each of these mutant strains (both strains of mice were homozygous for their respective mutations/transgenes), as well as wild type *APP*$^{+/+}$ and non-transgenic (non-Tg) controls, were subjected to microarray analysis at 12 months of age using the Affymetrix Mouse Gene 2.0ST Array. All mice used in the study were maintained as inbred lines. There is no information on whether any of the mice analysed were littermates. It is highly unlikely that the mice used in each comparison between mutant individuals and their wild type counterparts all arose from the same litter, because obtaining 3 homozygous and 3 wild type male mice in a single litter arising from an in-crossing of heterozygous mutant mice, (expected to produce a wild type : heterozygote : homozygote Mendelian genotype ratio of 1:2:1), would be a rare event as litters of mice generally consist of 5 to 10 pups. Therefore, additional variation was introduced into the analysis through use of mice from different litters and this is likely confounding with genotype. This is important to note, as the results presented here were generated under the assumption that any effects of litter of origin are negligible.

We first obtained the raw microarray data from the GEO database (accession number GSE92926). Initially, we attempted to replicate the results of Castillo et al. (2017) using the Affymetrix Transcriptome Analysis Console software. However, we were unable to find sufficient information to replicate their results. Therefore, we analysed the microarray dataset in a reproducible manner by importing the .CEL files for all twelve mice for analysis with *R* (Team, 2019) using the *oligo* package (Carvalho and Irizarry, 2010). The analysis of this dataset is based on the proposed workflow found in (Klaus and Reisenauer, 2018) and code to reproduce the analysis can be found at https://github.com/karissa-b/AD-signature/.

### Pre-processing of raw data

Raw intensities were pre-processed using the robust multichip average (rma) method (Irizarry et al., 2003). We next excluded probesets which contained a median log2 intensity value of < 3.5 (lowly expressed) and also any probesets assigned to multiple genes as recommended by Klaus and Reisenauer (2018). The distribution of the intensity values before and after the pre-processing can be found in **Figure 1**.

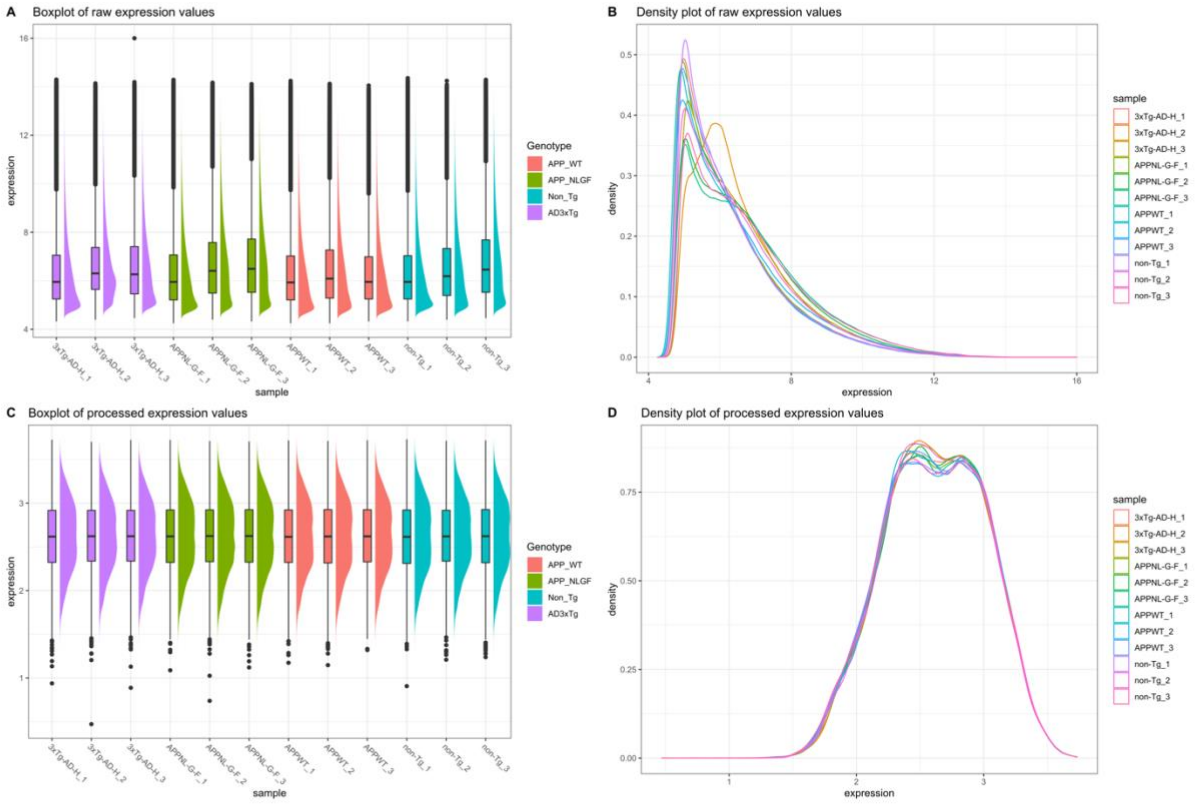


**Figure 1: A)** Boxplot and **B)** density plot of the raw intensity data. **C)** Boxplot and **D)** density plot of the intensity data after rma normalisation and filtering for lowly expressed and multi-mapping probes.

### Principal component analysis

We performed principal component analysis (PCA) to explore the overall similarity between samples. **Figure 2** below shows the plot of principal component 1 (PC1) against PC2 for each microarray sample after pre-processing. Samples separated across PC1 by genotype, suggesting that the homozygous genotypes in this study result in distinct transcriptome states. Notably, the $App^{NL\text{-}G\text{-}F/NL\text{-}G\text{-}F}$ samples and their corresponding $App^{+/+}$ control samples appear to separate to a greater extent across PC1 than the 3xTg samples and their corresponding non-Tg wild type control samples. This suggests that the disturbance to the cortex transcriptome in $App^{NL\text{-}G\text{-}F/NL\text{-}G\text{-}F}$ mice is greater than that in 3xTg mice.

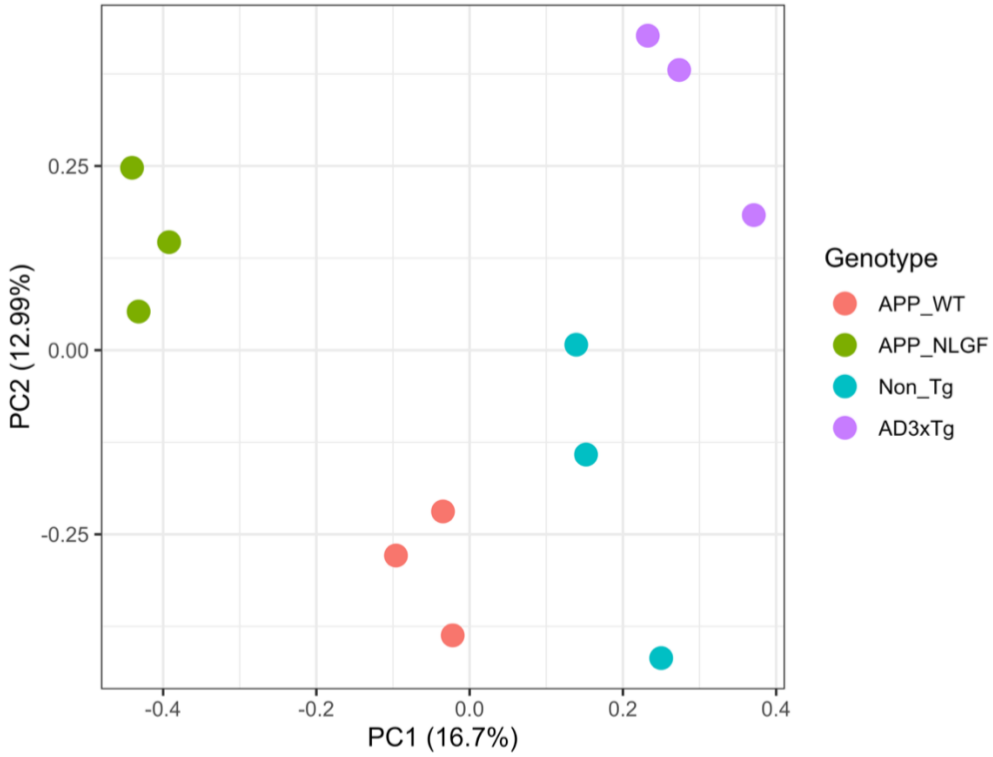


**Figure 2:** Principal component (PC) analysis on rma-processed intensity values.

### Differential Gene Expression Analysis

To determine which probesets (i.e. genes) are differentially expressed (DE) in each of the pairwise comparisons of mutant mice with their respective non-mutant/non-Tg controls, we performed differential gene expression analysis with *limma* (Ritchie et al., 2015). A design matrix was generated to specify sample genotypes, while a contrasts matrix was generated to specify the pairwise contrasts. We considered a gene to be DE if the FDR-adjusted p-value was less than 0.05 (no fold change filter). We identified 158 genes to be differentially expressed (DE) in *App*$^{NL\text{-}G\text{-}F/NL\text{-}G\text{-}F}$ mice relative to *App*$^{+/+}$ mice, and 126 genes to be DE in 3xTg mice relative to non-Tg wild type controls (**Figure 3A**, **B**). Three downregulated genes: *Fos*, *Gadd45b,* and *Nr4a1* and one upregulated gene, *Il33,* were found to be DE in both forms of mutant mice relative to their wild type/non-transgenic counterparts (**Figure 3C, D**).

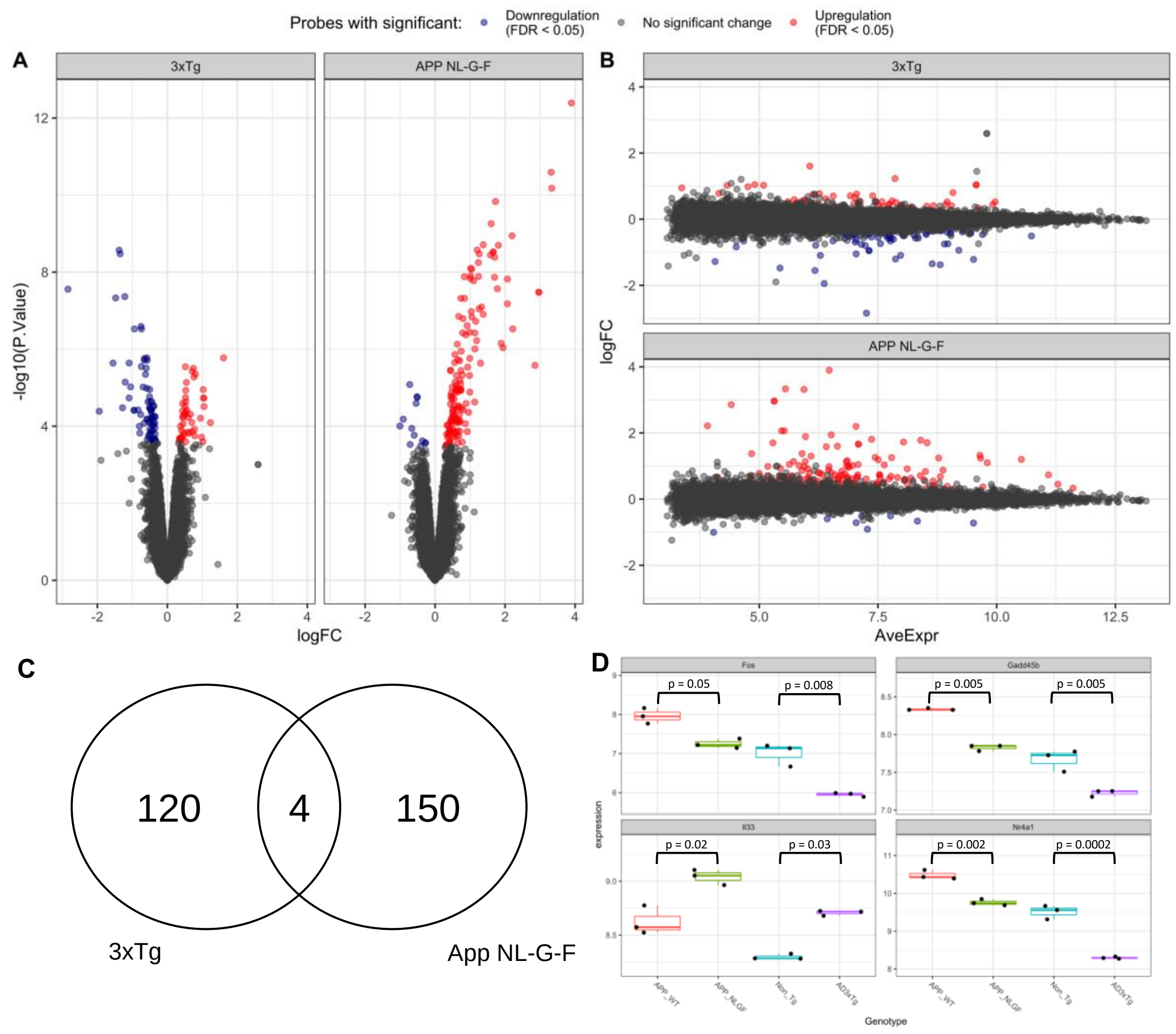


**Figure 3:** Differential gene expression analysis. **A)** Volcano plot and **B)** MD plot of the changes to gene expression in *App*$^{NL\text{-}G\text{-}F/NL\text{-}G\text{-}F}$ and 3xTg mice relative to controls. **C)** Venn diagram showing four genes are identified as differentially expressed (DE) in both comparisons. **D)** Boxplot of expression values of the four shared DE genes. The FDR-adjusted p-values as calculated from the *limma* analysis are indicated.

### Over-representation analysis

We next tested whether any *KEGG* (Kanehisa and Goto, 2000) or *IRE* (Hin et al., 2020) gene sets were significantly over-represented within the DE genes in each comparison using *kegga* (Young et al., 2010). The *KEGG* gene sets were obtained using the package *msigdbr*

(Dolgalev, 2020), and the *IRE* (Hin et al., 2020) gene sets, obtained from https://github.com/nhihin/ire. We restricted the gene sets to include only genes that had been tested for differential expression in the *limma* analysis. After correcting for multiple testing using the Bonferroni method, we found statistical evidence (Bonferroni-adjusted p-value < 0.05) for two gene sets as significantly over-represented among the DE genes in 3xTg mice, and 27 gene sets to be significantly overrepresented among the DE genes in *App*$^{NL\text{-}G\text{-}F/NL\text{-}G\text{-}F}$ mice (**Figure 4A**). No *IRE* gene sets were found to be significantly enriched among either of the DE gene lists. Notably, the enriched DE genes within each gene set in the *App*$^{NL\text{-}G\text{-}F/NL\text{-}G\text{-}F}$ mice appear to be shared across many of the gene sets (**Figure 4B**). Many of these gene sets contain shared DE genes are involved in inflammatory processes, which is not unexpected as gliosis has previously been observed to occur in these mutant mice (Castillo et al., 2017; Saito et al., 2014). Only the *KEGG_LYSOSOME* gene set appears to be enriched in mostly independent DE genes (which are upregulated) (**Figure 4C**). In summary, enrichment testing within the DE genes suggests that, at 12 months of age, the *App*$^{NL\text{-}G\text{-}F/NL\text{-}G\text{-}F}$ mice have a more severe phenotype/transcriptome disturbance than the 3xTg mice (due to the larger number of gene sets significantly altered), and this phenotype mostly consists of an inflammatory response, and changes to the lysosome.

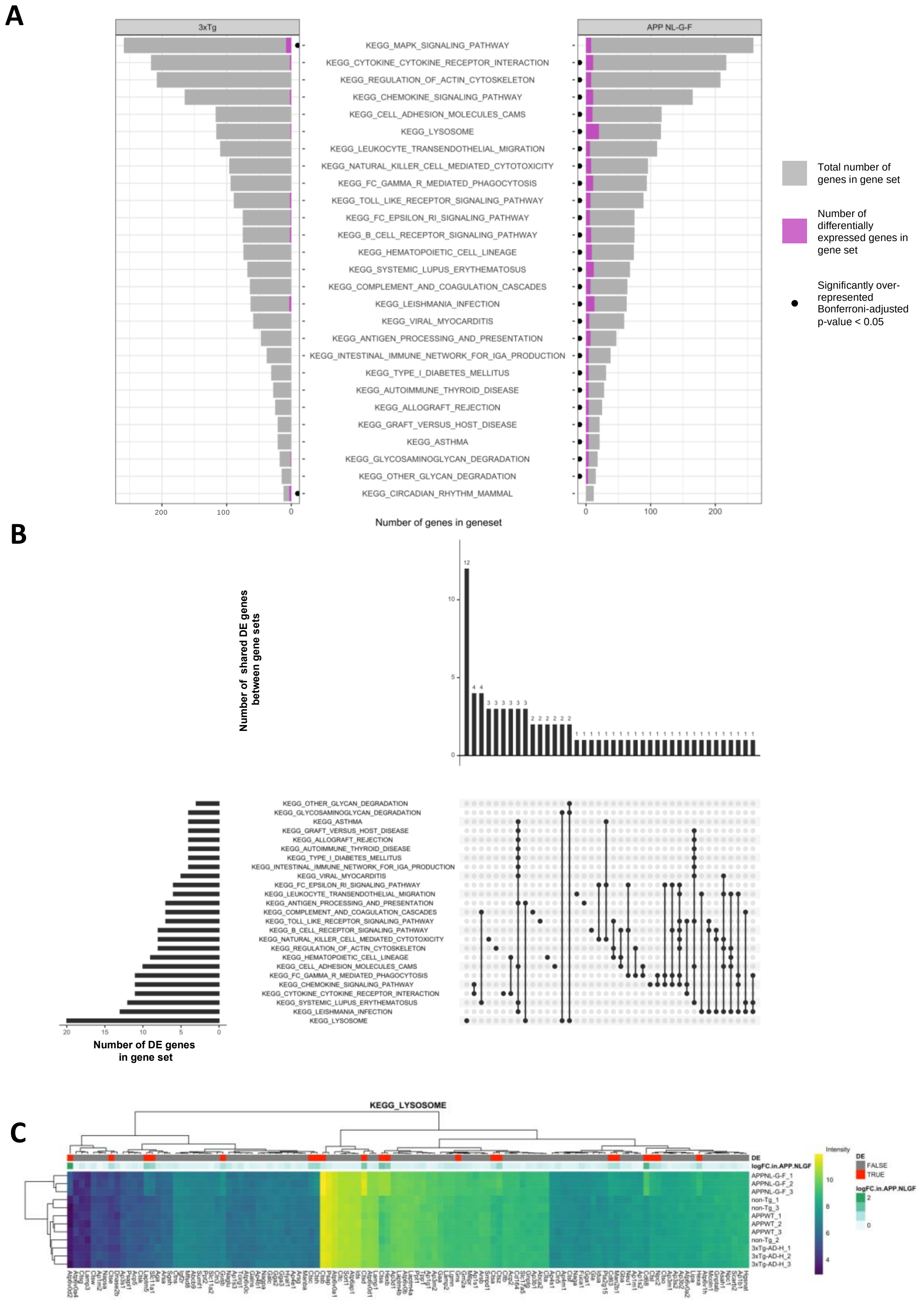
A
3xTg
APP NL-G-F
KEGG_MAPK_SIGNALING_PATHWAY
KEGG_CYTOKINE_CYTOKINE_RECEPTOR_INTERACTION
KEGG_REGULATION_OF_ACTIN_CYTOSKELETON
KEGG_CHEMOKINE_SIGNALING_PATHWAY
KEGG_CELL_ADHESION_MOLECULES_CAMS
KEGG_LYSOSOME
KEGG_LEUKOCYTE_TRANSENDOTHELIAL_MIGRATION
KEGG_NATURAL_KILLER_CELL_MEDIATED_CYTOTOXICITY
KEGG_FC_GAMMA_R_MEDIATED_PHAGOCYTOSIS
KEGG_TOLL_LIKE_RECEPTOR_SIGNALING_PATHWAY
KEGG_FC_EPSILON_RI_SIGNALING_PATHWAY
KEGG_B_CELL_RECEPTOR_SIGNALING_PATHWAY
KEGG_HEMATOPOIETIC_CELL_LINEAGE
KEGG_SYSTEMIC_LUPUS_ERYTHEMATOSUS
KEGG_COMPLEMENT_AND_COAGULATION_CASCADES
KEGG_LEISHMANIA_INFECTION
KEGG_VIRAL_MYOCARDITIS
KEGG_ANTIGEN_PROCESSING_AND_PRESENTATION
KEGG_INTESTINAL_IMMUNE_NETWORK_FOR_IGA_PRODUCTION
KEGG_TYPE_I_DIABETES_MELLITUS
KEGG_AUTOIMMUNE_THYROID_DISEASE
KEGG_ALLOGRAFT_REJECTION
KEGG_GRAFT_VERSUS_HOST_DISEASE
KEGG_ASTHMA
KEGG_GLYCOSAMINOGLYCAN_DEGRADATION
KEGG_OTHER_GLYCAN_DEGRADATION
KEGG_CIRCADIAN_RHYTHM_MAMMAL
200
100
0
Number of genes in geneset
Total number of genes in gene set
Number of differentially expressed genes in gene set
Significantly over-represented Bonferroni-adjusted p-value < 0.05
B
Number of shared DE genes between gene sets
Number of DE genes in gene set
C
KEGG_LYSOSOME
DE
logFC.in.APP.NLGF
FALSE
TRUE
Intensity
APPNL-G-F_1
APPNL-G-F_2
APPNL-G-F_3
non-Tg_1
non-Tg_3
APPWT_1
APPWT_2
APPWT_3
non-Tg_2
3xTg-AD-H_1
3xTg-AD-H_2
3xTg-AD-H_3

**Figure 4**: Over-representation analysis. **A)** Pyramid bar plot indicating the number of genes in the significantly enriched KEGG and IRE gene sets in *App*$^{NL\text{-}G\text{-}F/NL\text{-}G\text{-}F}$ and 3xTg mice. Only gene sets with a Bonferroni adjusted p-value from *kegga* are shown (and are indicated by a black dot). The total numbers of genes in these gene sets are shown by grey bars, while the numbers of significantly differentially expressed genes in these gene sets are shown in magenta. **B)** Upset plot indicating the high degree of overlap of DE genes across the significantly enriched gene sets in *App*$^{NL\text{-}G\text{-}F/NL\text{-}G\text{-}F}$ mice. **C)** Heatmap of the *KEGG_LYSOSOME* gene set indicating the intensity expression values. Both rows and columns are clustered according to their Euclidean distance. Genes are labelled red if they were found to be differentially expressed by *limma* in *App*$^{NL\text{-}G\text{-}F/NL\text{-}G\text{-}F}$ mice, and the magnitude of logFC is shown in green.

### Ranked list enrichment analysis

We next obtained a more complete view on the changes to gene expression observed in *App*$^{NL\text{-}G\text{-}F/NL\text{-}G\text{-}F}$ and 3xTg mice relative to their respective controls by performing three methods of rank-based, gene set enrichment analysis: *fry* (Wu et al., 2010), *camera* (Wu and Smyth, 2012) and *GSEA* (Sergushichev, 2016; Subramanian et al., 2005). We then calculated the harmonic mean p-value (Wilson, 2019) from the raw p-values from each method as discussed in (Barthelson et al., 2020). In this analysis, we considered a *KEGG* or *IRE* gene set to be significantly altered if the Bonferroni-adjusted harmonic mean p-value was less 0.05. We observed similar gene sets to be significantly altered using this ranked-list approach to the over-representation analysis described above. Additionally, the significance of the gene sets is likely being driven by similar genes, as supported by the observed number of the DE genes across the gene sets (and to a lesser extent, the leading edge genes from the *GSEA* algorithm) (**Figure 5**). A summary of whether a gene set is observed to be significantly altered in either over-representation analysis or from our ranked-list approach can be found in **Figure 6**.

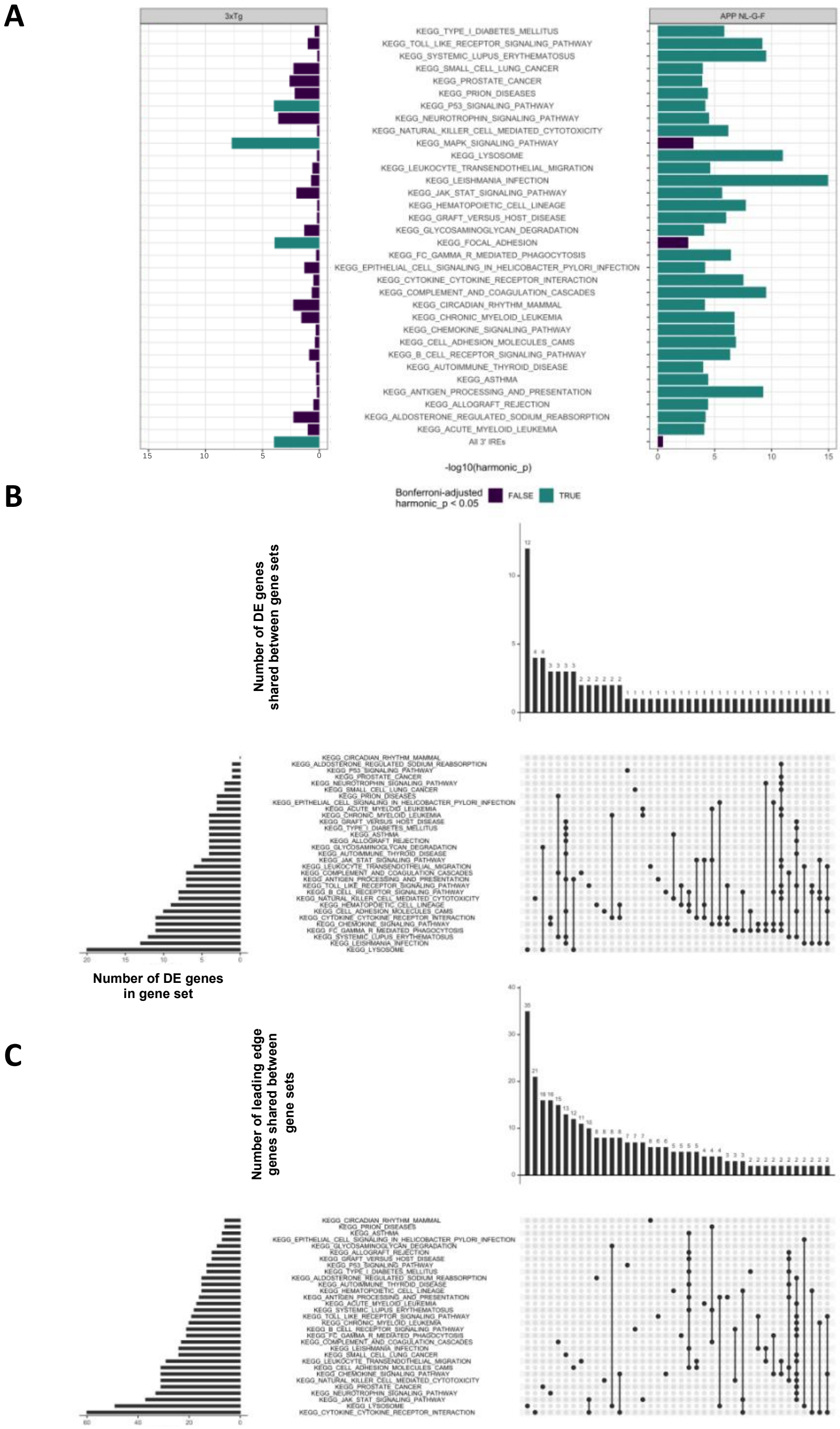


**Figure 5:** Ranked-list enrichment testing. **A)** Summary of significantly enriched *KEGG* and *IRE* gene sets in $App^{NL\text{-}G\text{-}F/NL\text{-}G\text{-}F}$ and 3xTg mice. Gene sets are coloured according to whether they were below the threshold of a Bonferroni-adjusted harmonic mean p-value of < 0.05. **B)** Upset plot indicating the overlap of DE genes across the significantly altered gene sets in $App^{NL\text{-}G\text{-}F/NL\text{-}G\text{-}F}$ mice. **C)** Upset plot indicating the leading edge genes from the *GSEA* algorithm in $App^{NL\text{-}G\text{-}F/NL\text{-}G\text{-}F}$ mice.

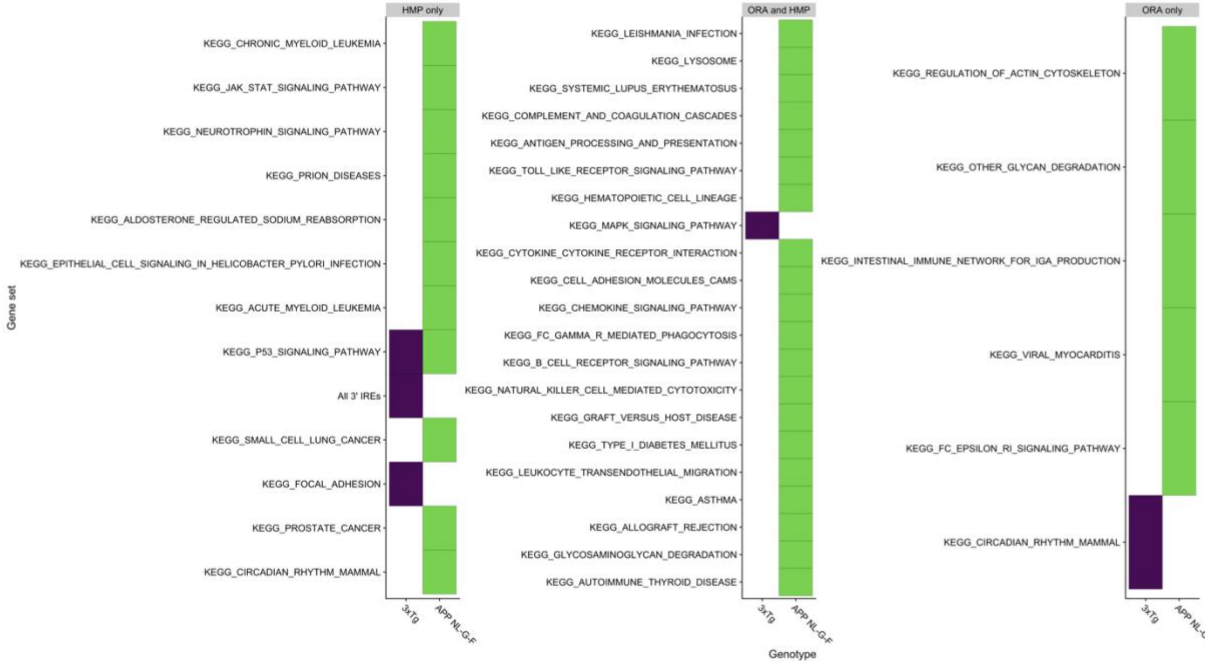


**Figure 6:** Summary of significantly altered gene sets in *App*$^{NL\text{-}G\text{-}F/NL\text{-}G\text{-}F}$ and 3xTg mice. Gene sets only found to be altered by calculation of the harmonic mean p-value (HMP) are shown on the left. Gene sets only found to be altered by over-representation analysis (ORA) using *kegga* are shown on the right. Gene sets found to be altered in both types of enrichment are shown in the middle. Gene sets are coloured according to the comparison in which they are significantly altered.

### Changes to cell type proportions are present in *App*$^{NL\text{-}G\text{-}F/NL\text{-}G\text{-}F}$ mice

Castillo et al. (2017) noted an increase in expression of marker genes of astrocytes, microglia and oligodendrocytes, while relatively consistent expression of marker genes of neurons in *App*$^{NL\text{-}G\text{-}F/NL\text{-}G\text{-}F}$ mice. However, only a maximum of 16 genes were used as markers of these cell types. Therefore, we inspected larger sets of genes obtained from (Cahoy et al., 2008) and (Oosterhof et al., 2017) to assess whether cell type proportions are altered in these microarray samples. Using *fry* with a directional hypothesis, we observed statistical evidence for increased expression of marker genes of astrocytes, oligodendrocytes and microglia (FDR < 0.05) in *App*$^{NL\text{-}G\text{-}F/NL\text{-}G\text{-}F}$ mice, suggesting increased abundances of these broad cell types found in the mouse brain (**Figure 7**). Approximately half of the DE genes in the *App*$^{NL\text{-}G\text{-}F/NL\text{-}G\text{-}F}$ mice overlap with marker genes of microglia.

Therefore, a significant proportion of the changes to gene expression we observed in this analysis may be artefactual due to increased proportions of microglia in the *App*$^{NL\text{-}G\text{-}F/NL\text{-}G\text{-}F}$ samples.

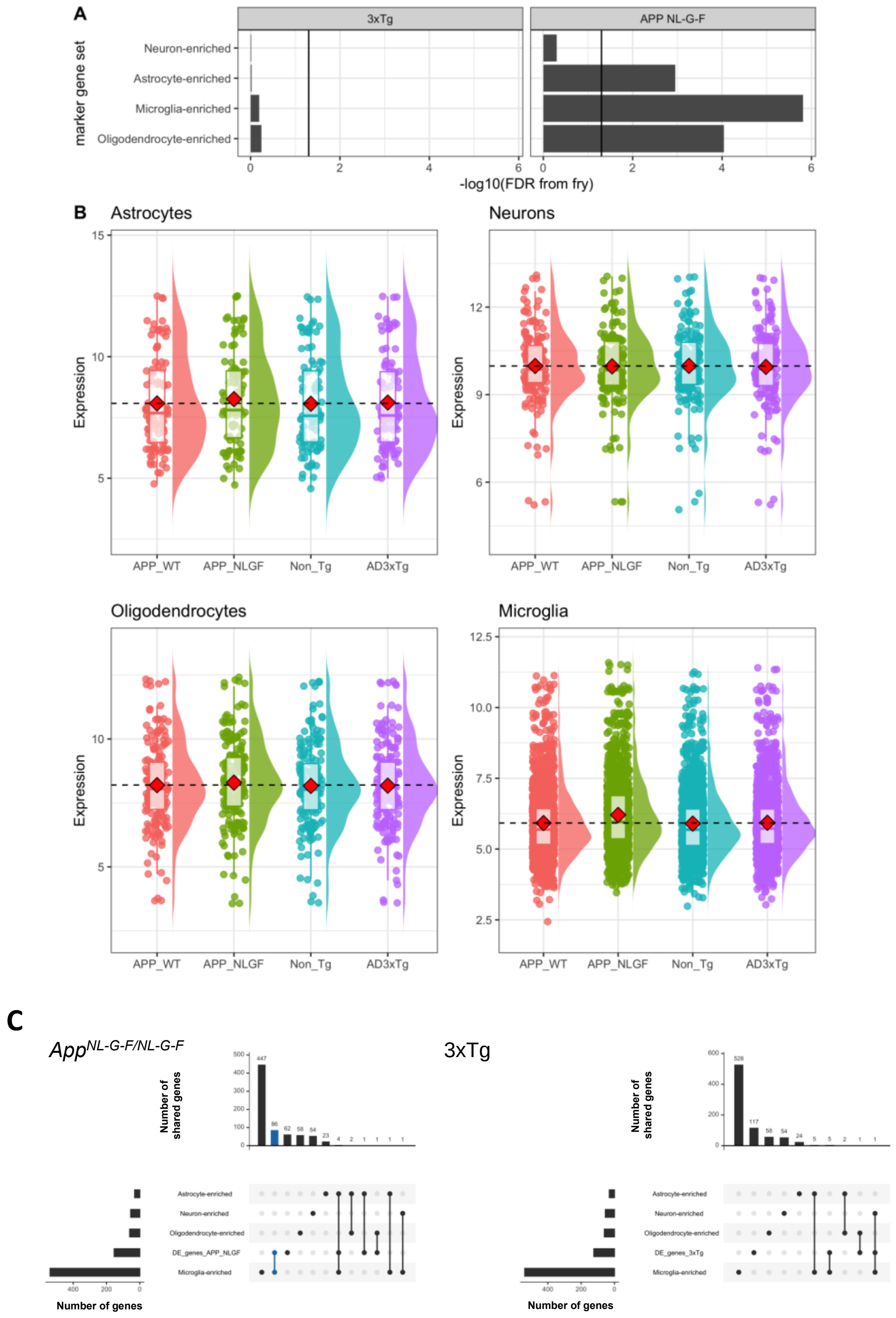

**Figure 7:** Proportions of cell types in *App*$^{NL-G-F/NL-G-F}$ mice are altered. **A)** Gene set testing using *fry* on marker gene sets of neurons, astrocytes and oligodendrocytes from (Cahoy et al., 2008) and microglia from (Oosterhof et al., 2017). The black line indicates an FDR-adjusted p-value of 0.05. **B)** Distribution of intensities of the marker genes across genotypes. The boxplots indicate summary statistics. The mean intensity value for each genotype is indicated by the red diamonds. To assist with visualisation of the increased expression of marker genes in *App*$^{NL-G-F/NL-G-F}$ mice, the mean expression in *App*$^{+/+}$ mice is also shown as a black dashed line. **C)** Upset plot showing the overlap of DE genes found in *App*$^{NL-G-F/NL-G-F}$ (left) and 3xTg (right) mice with the cell type marker gene sets.

### Discussion and Conclusion

In summary, we draw three main conclusions from our re-analysis of the microarray transcriptomic data from the cortices of 12 month old male *App*$^{NL-G-F/NL-G-F}$ mice and 3xTg mice:

1) *App*$^{NL-G-F/NL-G-F}$ mice and 3xTg mice show distinct brain transcriptomic disturbances relative to controls.
2) Brain transcriptomes from *App*$^{NL-G-F/NL-G-F}$ mice, when compared to transcriptomes wild type mice, show apparent strong inflammatory signals and changes to gene expression implicating the lysosome.
3) A significant factor contributing (artefactually) to apparent disturbances of gene expression observed in *App*$^{NL-G-F/NL-G-F}$ mice is likely increased proportions of microglia within the cortex samples

Our re-analysis of the microarray dataset described by Castillo et al. (2017) mostly replicated their conclusions. At the single gene level, we found some overlap of DE genes with Castillo et al. (2017), and some distinct DE genes (data not shown). This is likely due to different approaches used in pre-processing of the raw data and testing for differential gene expression. However, at the pathway (i.e. gene set) level, both analyses predict similar processes to be affected in each mouse model.

We also observed highly significant upregulation of genes involved in lysosome function in *App*$^{NL\text{-}G\text{-}F/NL\text{-}G\text{-}F}$ mice but not in 3xTg mice. Changes to the lysosomal network have been observed previously in *App*$^{NL\text{-}G\text{-}F/NL\text{-}G\text{-}F}$ mice at a young age at the protein level (Whyte et al., 2020), supporting the validity of this result. The Iberian mutation of *APP* present in *App*$^{NL\text{-}G\text{-}F/NL\text{-}G\text{-}F}$ mice has been shown to increase levels of the C99 fragment of APP (Guardia-Laguarta et al., 2010), which would impair acidification of the endo-lysosomal system (Jiang et al., 2019), and likely result in an intracellular ferrous iron deficiency, mitochondrial dysfunction and inflammation (Yambire et al., 2019). We did not observe any evidence for mitochondrial dysfunction or iron dyshomeostasis in this dataset. However, we suspect that these signals may be lost in the noise due to sample litter-of-origin issues and the strong inflammatory signal.

From this dataset, it appears that the *App*$^{NL\text{-}G\text{-}F/NL\text{-}G\text{-}F}$ mice have a more severe phenotype than the 3xTg mouse at 12 months of age. This is unexpected, as the more extensive genomic differences of the 3xTg mice relative to *App*$^{NL\text{-}G\text{-}F/NL\text{-}G\text{-}F}$ mice, would, in a simplistic view, be thought to affect more cellular processes. Indeed, 3xTg mice show cognitive impairment from 4 months age (Billings et al., 2005), while *App*$^{NL\text{-}G\text{-}F/NL\text{-}G\text{-}F}$ mice do not show cognitive impairment until 6 months of age (Mehla et al., 2019). However, the memory impairments at 12 months of age appeared to be most severe in *App*$^{NL\text{-}G\text{-}F/NL\text{-}G\text{-}F}$ mice (Mehla et al., 2019), which would imply that the effects on cellular processes are also more severe at this age.

Notably, the comparisons made in this microarray analysis between the *App*$^{NL\text{-}G\text{-}F/NL\text{-}G\text{-}F}$ and *App*$^{+/+}$ mice observed the compounded effects of homozygosity for 6 mutations simultaneously. An interesting future experiment could entail comparison of the *App*$^{NL\text{-}G\text{-}F/NL\text{-}G\text{-}F}$ mice to mice carrying only the humanised Aβ sequence (i.e. (Serneels et al., 2020)) to assist with identification of changes to the transcriptome due specifically to the EOfAD mutations in this knock-in mouse model. Finally, the inclusion of three EOfAD mutations in *APP* within the same animal is of questionable relevance to the genetic state of human EOfAD. Therefore, the results presented here are not ideal for comparison with the transcriptome changes observed in our heterozygous, single knock-in mutation zebrafish models. The $App^{NL}$ mouse model carries only the Swedish *APP* EOfAD mutation within the humanised Aβ sequence (Saito et al., 2014). A brain transcriptome analysis of young (3 or 6 months of age) heterozygous $App^{NL}$ mice, relative to mice carrying only the humanised Aβ sequence, would not require the generation of new mouse mutants and would provide data more comparable with that already available from our studies using zebrafish.
