## Supplemental File 3 for "Comparative analysis of Alzheimer’s disease knock-in model brain transcriptomes implies changes to energy metabolism as a causative pathogenic stress"

Possession of the ε4 allele of apolipoprotein E (*APOE*) is the greatest genetic risk factor for development of sporadic, late-onset Alzheimer's disease (LOAD). Homozygosity for the ε4 allele increases an individual's risk at least 9-fold relative to the common, ε3 allele (depending on sex and ethnicity) (Farrer et al., 1997). There is also an ε2 allele of *APOE*, which is protective against development of AD (Farrer et al., 1997). To understand the effects of the different alleles of *APOE*, targeted replacement mice have been developed expressing humanised ε2, ε3, or ε4 alleles from the mouse *Apoe* locus (APOE-TR) (Sullivan et al., 1997).

Zhao et al. (2020) performed an RNA-seq experiment investigating the effect of *APOE* genotype, age (3, 12 and 24 months of age) and sex in APOE-TR homozygous mice. In that analysis, pairwise comparisons between the ε2, or ε4 alleles relative to the ε3 allele were not conducted at each age and sex. Only genes/pathways which were influenced overall by *APOE* genotype, age, sex , and interactions between these factors was reported. We are interested in the cellular processes implicated as affected by AD-causative mutations in young brains. Therefore, here we will re-analyse the RNA-seq dataset from Zhao et al. to ask which processes are affected by homozygosity for the ε2, or ε4 alleles relative to the ε3 allele of *APOE* in the brains of three and twleve month old mice. Throughout this analysis, we refer to these homozygous mice as "APOE2", "APOE3" and "APOE4".

### RNA-seq data processing

We first obtained the raw fastq files for the entire APOE-TR RNA-seq experiment from AD Knowledge Portal (accession number syn20808171, https://adknowledgeportal.synapse.org/). The raw reads were first processed using *AdapterRemoval* (version 2.2.1) (Schubert et al., 2016), setting the following options: --*trimns*, *--trimqualities, --minquality 30, --minlength 35.* Then, the trimmed reads were aligned to the *Mus musculus* genome (Ensembl GRCm38, release 98) using *STAR* (version 2.7.0) (Dobin et al., 2013) using default parameters. The gene expression counts matrix was generated using *featureCounts* (version 1.5.2) (Liao et al., 2014). We only counted the number of reads which uniquely aligned to, strictly, exons with a mapping quality of at least 10 to predict expression levels of genes in each sample.

We then imported the output from *featureCounts* (Liao et al., 2014) for analysis with *R* (Team, 2019). We first omitted genes which are lowly expressed (and are uninformative for differential expression analysis). We considered a gene to be lowly expressed if it contained, at most, 2 counts per million (CPM) in 8 or more of the 24 samples we analysed. The effect of filtering lowly expressed genes is found in **Figure 1** below.

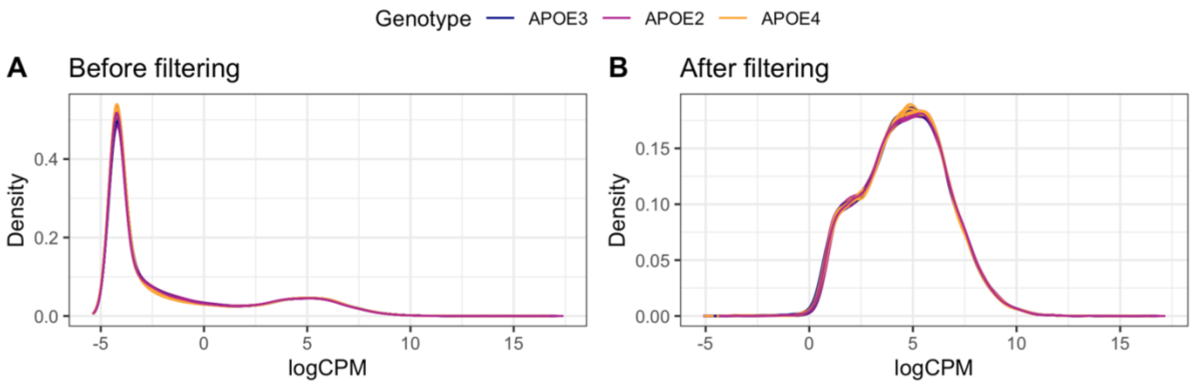


**Figure 1:** The density of the log counts per million (logCPM) values detected in 3 month old mouse brain samples is shown before filtering in **A**, then after omitting samples with a logCPM of < 2 in at least one third of the samples in **B**.

### Assessment of expression of genes from the Y-chromosome to confirm sex.

We then confirmed the sex of each mouse by examination of the expression of genes from the Y-chromosome. Sample APOE_3M_21 appears to be a male sample as it expresses genes from the Y chromosome. Samples  APOE_3M_30 and APOE_3M_7 appear to be female as they do not express genes from the Y-chromosome (**Figure 2**). This was subsequently corrected for the rest of the analysis.

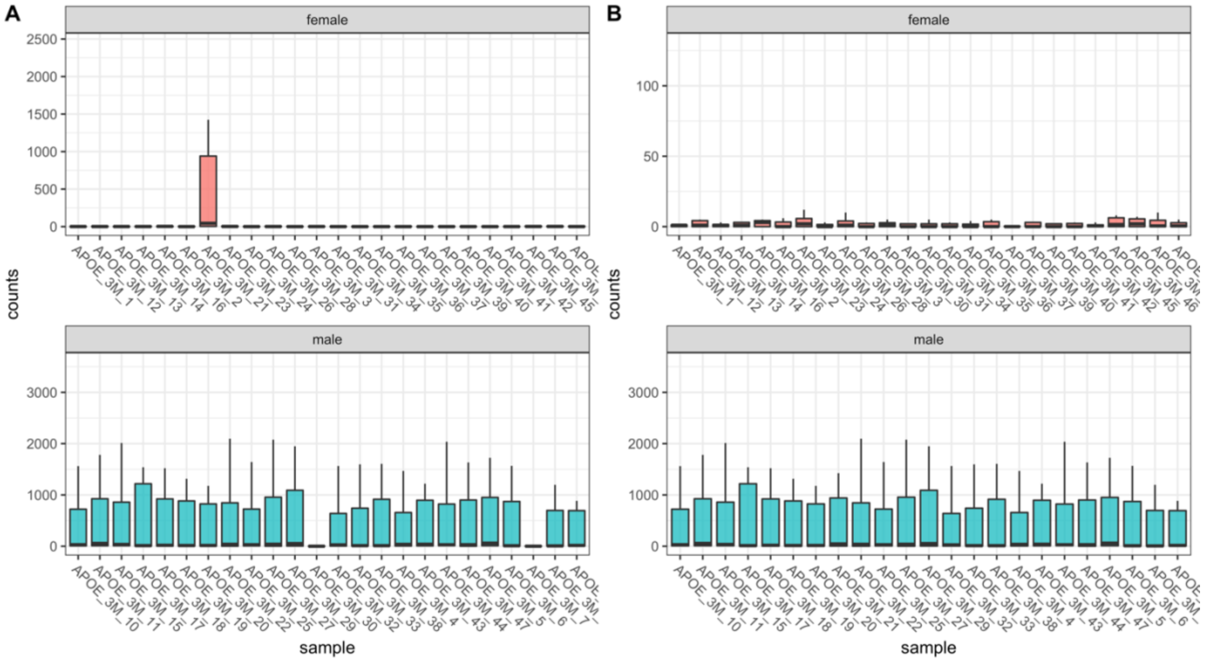


**Figure 2: A)** Boxplots showing the distribution of expression of male-specific genes (located on the Y-chromosome) in the cortex of 3 month old APOE-TR mice, grouped by initial sex. **B)** Expression of male-specific genes after correcting the sex of the samples.

### Principal component analysis

We next performed principal component analysis (PCA) to explore whether *APOE* genotype results in stark differences to the brain transcriptome at 3 months of age. A plot of principal component 1 (PC1) against principal component 2 (PC2) revealed that samples separated into two distinct clusters of sex across PC2 (**Figure 3A**). This suggests that the effect of sex on the murine brain transcriptome is substantial and cannot be ignored in the differential gene expression analysis. Among the male samples, APOE4 samples form a distinct cluster from the APOE2 and APOE3 samples, suggesting that the APOE4 genotype has a distinct effect on the transcriptome compared to APOE2 relative to APOE3 in males. This is not observed to the same extent in the female samples. However, the male APOE4 and APOE3 samples appeared to arise mostly from single litters, as implied from the date-of-birth of each sample (**Figure 3B**).

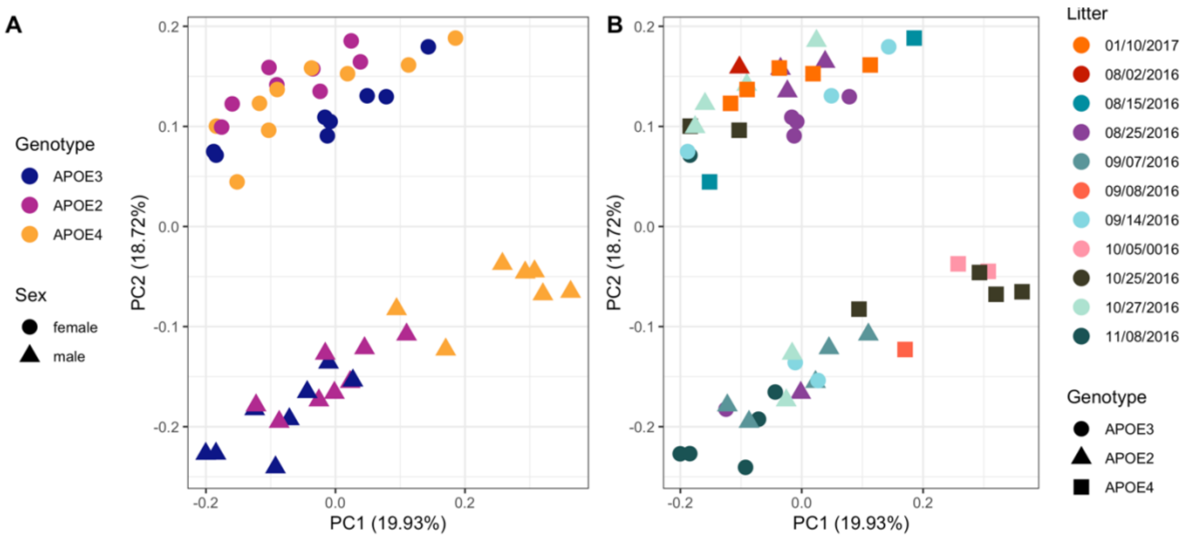


**Figure 3: Principal component analysis. A** shows principal component 1 (PC1) against PC2. The numbers between parentheses indicate the percentage of variation in the dataset that PC explains. Each point represents a sample, which is coloured by *APOE* genotype, and shaped by sex. **B** also shows PC1 against PC2. However, each point is coloured by litter (implied from the date-of-birth of each mouse), and shaped by *APOE* genotype.

From **Figure 3B**, we observed that some experimental groups of samples (i.e. the male APOE3 and male APOE4 samples) appeared to arise mostly from single litters of mice. This is confounding with the effect of genotype and complicates interpretation of whether any effects observed in a pairwise contrast between male APOE3 and APOE4 mice are due to APOE genotype or litter-of-origin (or, most likely, both). Indeed, $\chi 2$ tests for independence revealed that there is a highly significant dependence of *APOE* genotype and litter across the entire 3-month-old dataset ($\chi 2$= 82.7, df = 20, p-value = 1.4e-09), only within male samples ($\chi 2$= 43.1, df = 14, p-value = 8.2e-05) and only within female samples ($\chi 2$= 39.3, df = 14, p-value = 3.0e-04). Some litters did not contain sufficient mice to remove the effect (i.e. some coefficients could not be estimated during the generalised linear model fitting procedure due to the design matrix not having full rank). Therefore, we continued the analysis assuming that the effect of litter is negligible.

### Initial differential gene expression analysis

To determine which genes were dysregulated in APOE4 and APOE2 mice relative to APOE3, we performed a differential gene expression analysis using a generalised linear model and likelihood ratio tests using *edgeR* (McCarthy et al., 2012; Robinson et al., 2009). We chose a design matrix which specifies the *APOE* genotype and sex of each sample. The contrasts matrix was specified to compare the effect of APOE2 or APOE4 relative to APOE3 in both males and females. In this analysis, we considered a gene to be differentially expressed (DE) if the FDR-adjusted p-value was < 0.05. Many genes were found to be DE in each comparison, particularly in male APOE4 mice (**Figure 4A**). Additionally, the bias for GC content and gene length for differential expression noted by Zhao et al. in the original analysis was also apparent (**Figure 4A, B**).

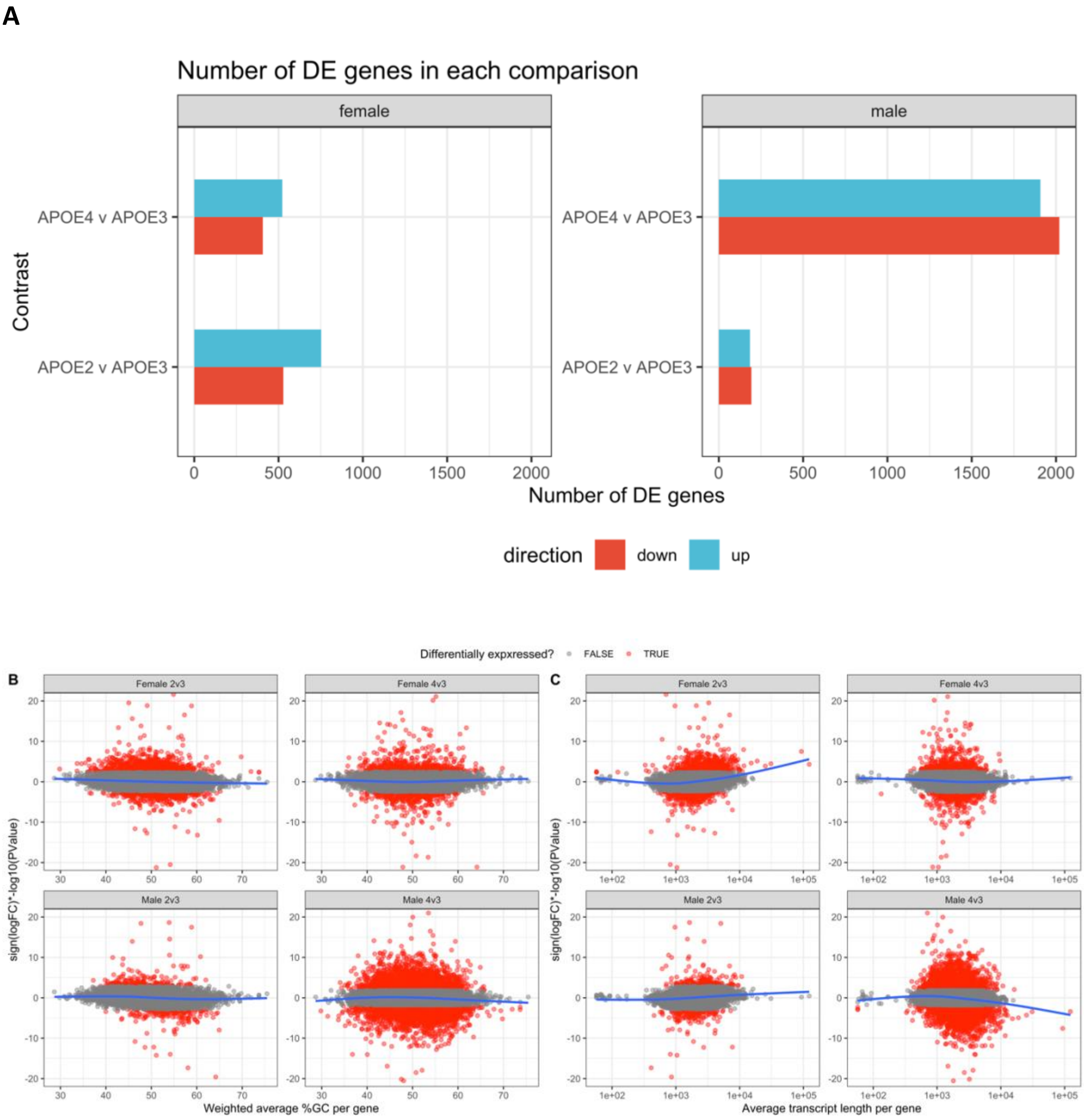


**Figure 4: Initial differential expression analysis. A)** Bar chart showing the number of differentially expressed genes (DE) in each comparison of the APOE4 or APOE2 genotype to APOE3. **B)** A ranking statistic per gene was calculated as the sign of the logFC multiplied by the negative log10 of the p-value from the likelihood ratio tests in *edgeR*. This was plotted against a weighted (by transcript length) average %GC content per gene and **C)** average transcript length. The blue generalised additive model fit (gam) lines are not centred on 0, indicating a bias.

### Conditional quantile normalisation

A gene length bias for differential expression has been shown to influence the results of gene set enrichment analyses (Mandelboum et al., 2019). Therefore, to correct for the observed bias for gene length (and GC content), we used conditional quantile normalisation (*cqn*) (Hansen et al., 2012). We calculated the average transcript length per gene, and a weighted (by transcript length) average %GC content per gene as input to *cqn* to produce the offset to correct for the bias. This offset was then included in an additional generalised linear model and likelihood ratio tests in *edgeR* with the same design and contrast matrices. In these tests, many genes were identified as DE and the bias for %GC and gene length had improved. A gene length bias was still present in male APOE4 and female APOE2 comparisons to APOE3. However, these were only observed due to a small number of long gene transcripts and likely can be ignored.

**A**

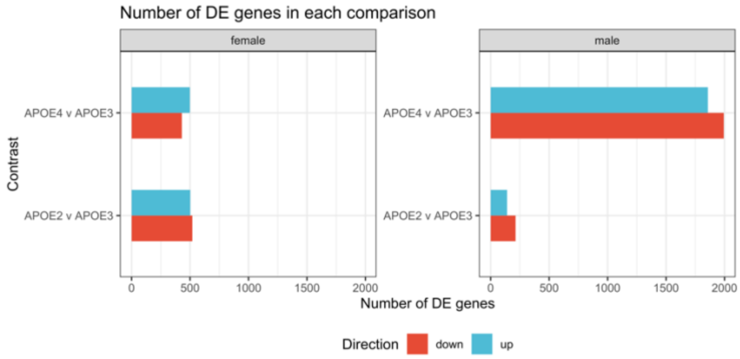

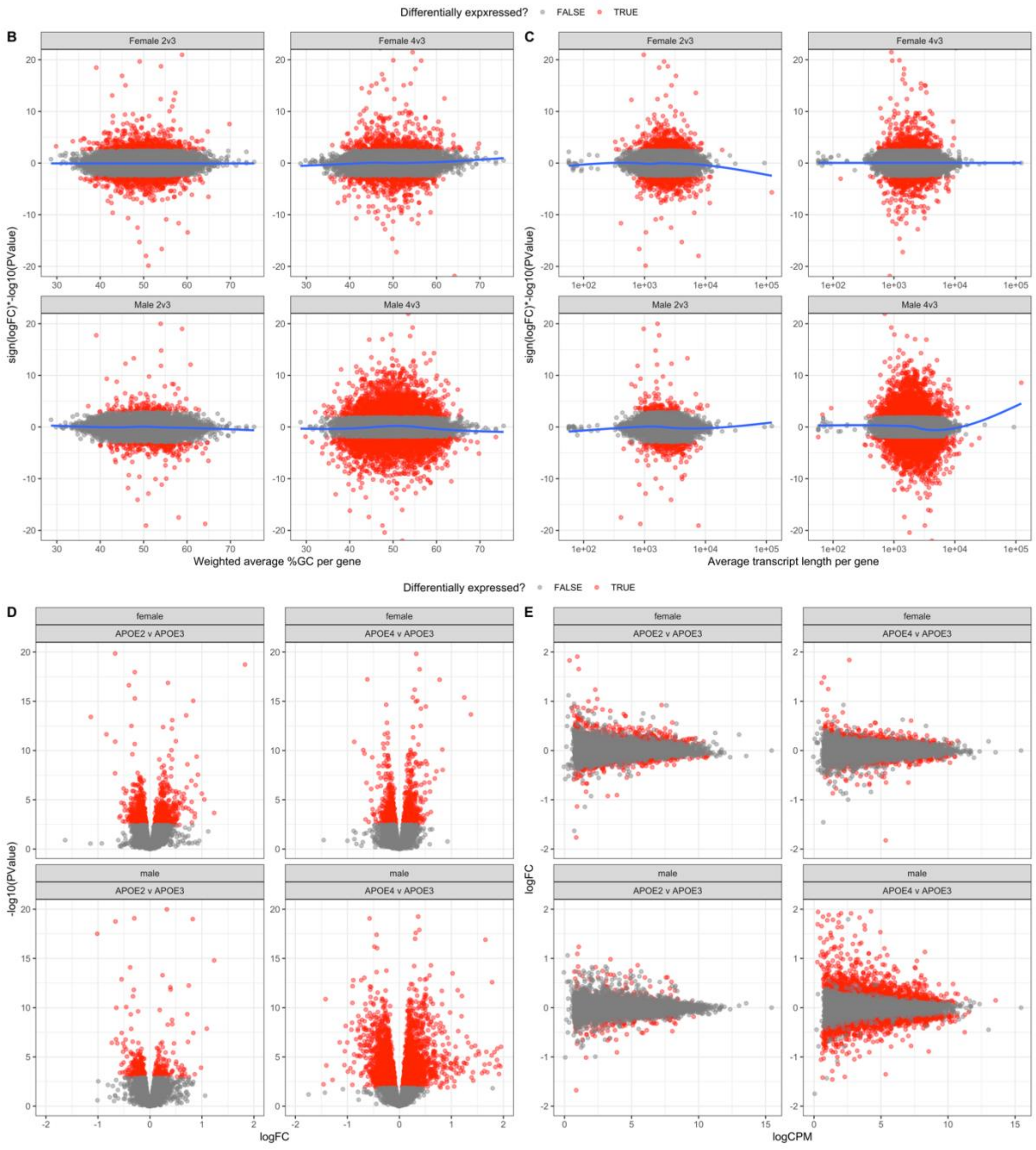


**Figure 5:** Differential gene expression analysis after *cqn*. **A)** Number of genes identified to be differentially expressed (DE) after *cqn*. **B)** Improvement of observed bias between %GC content and **C)** gene length for differential expression after *cqn*. The remaining bias for transcript length in the female APOE2 and male APOE4 comparisons appear to be only driven by a small number of genes. **D)** Volcano plots and **E)** mean difference (MD) plots of the changes to gene expression observed due to homozygosity for the APOE4 or APOE2 alleles relative to APOE3 in male and female mice. The limits of the x-axis in **D)** and the y-axis in **E)** are constrained to -2 and 2, and of the y-axis in **D)** to between 0 and 20, for visualisation purposes.

### Enrichment analysis

We next tested for over-representation of the *KEGG* (Kanehisa and Goto, 2000) and *IRE* (Hin et al., 2020) gene sets within the DE gene lists using *goseq* (Young et al., 2010), using the average transcript length per gene to calculate the probability weighting function (PWF). After correction for multiple testing using the FDR, we observed a total of three gene sets to be significantly enriched in any of the DE gene lists. Two of these *KEGG* gene sets were enriched in the DE genes for male APOE4 mice: *KEGG_CALCIUM_SIGNALING_PATHWAY* ($p_{FDR}$ = 0.03) and *KEGG_NEUROACTIVE_LIGAND_RECEPTOR_INTERACTION* ($p_{FDR}$ = 0.04). Approximately one third of the DE genes in each of these gene sets are shared, indicating that the enrichments of these gene sets are driven partially by the same genes (**Figure 6**).

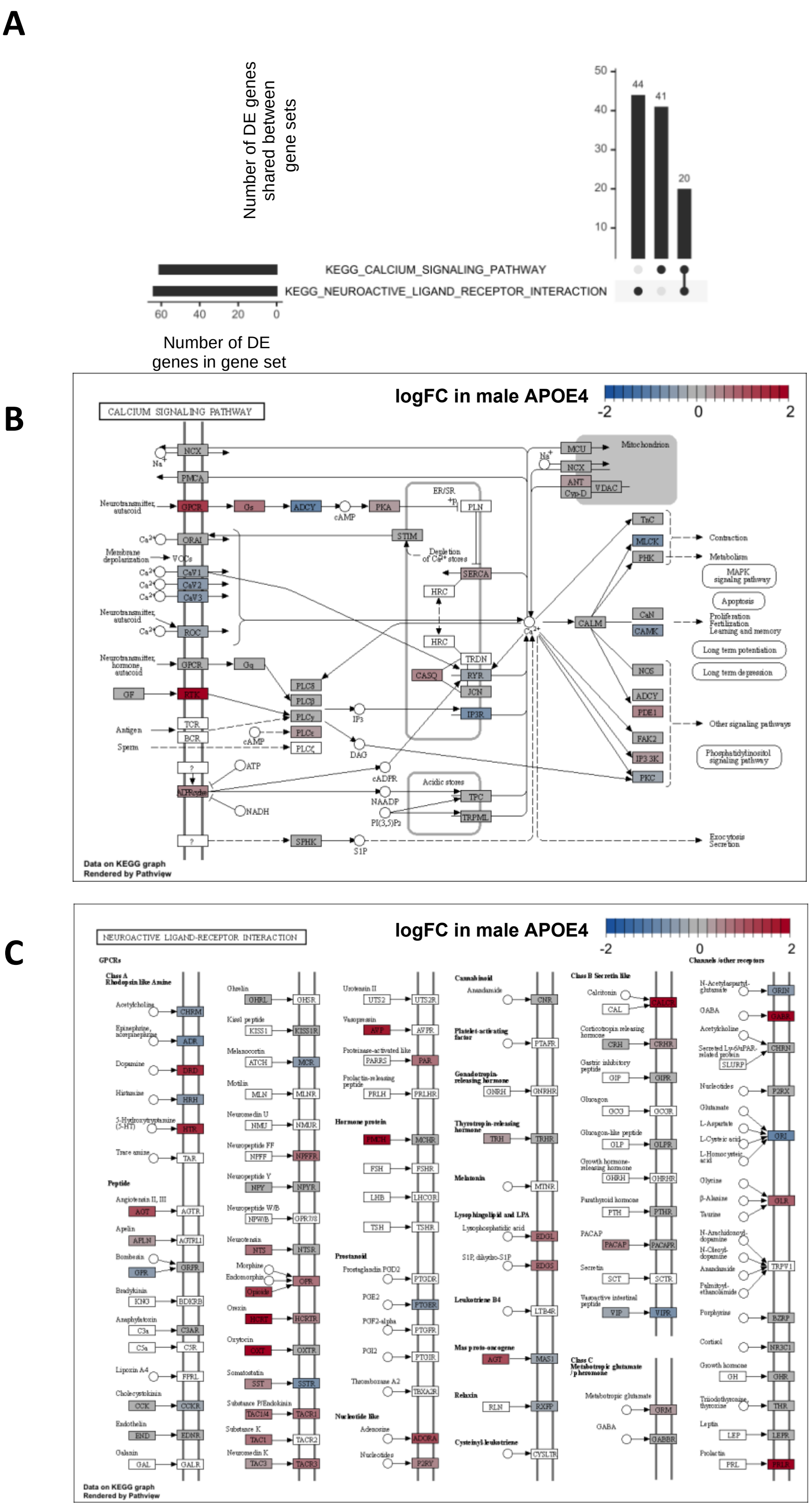

A
Number of DE genes shared between gene sets
50
40
30
20
10
44
41
20
KEGG_CALCIUM_SIGNALING_PATHWAY
KEGG_NEUROACTIVE_LIGAND_RECEPTOR_INTERACTION
60
40
20
0
Number of DE genes in gene set
B
CALCIUM SIGNALING PATHWAY
logFC in male APOE4
-2
0
2
Data on KEGG graph
Rendered by Pathview
C
NEUROACTIVE LIGAND-RECEPTOR INTERACTION
logFC in male APOE4
-2
0
2
Data on KEGG graph
Rendered by Pathview

**Figure 6:** Enrichment analysis within the lists of differentially expressed genes. **A)** Upset plot indicating the overlap of DE genes in male APOE4 samples for the significantly enriched gene sets. **B)** Pathview (Luo et al., 2017) visualisation of the logFC in male APOE4 samples for the *KEGG_CALCIUM_SIGNALING_PATHWAY* gene set and **C)** *KEGG_NEUROACTIVE_LIGAND_RECEPTOR_INTERACTION* gene set.

Additionally, the gene set *KEGG_STEROID_BIOSYNTHESIS* was found to be significantly enriched among the female APOE2 DE genes ($p_{FDR}$ = 1e-5). This gene set appears to be downregulated (**Figure 7**).

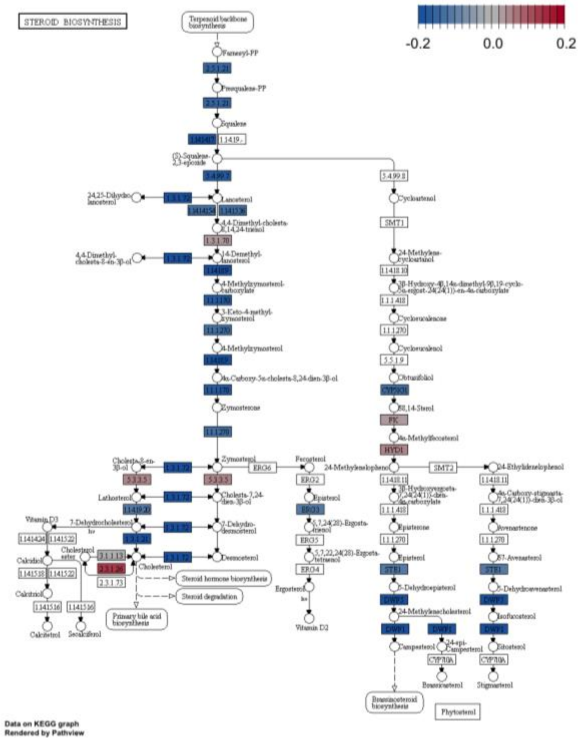


**Figure 7:** Pathview visualisation indicating the logFC of genes in the *KEGG_STEROID_BIOSYNTHESIS* gene set in female APOE2 mice.

Over-representation analysis using *goseq* relies on a hard threshold for a gene to be classified as DE. Therefore, information may be missed, i.e. genes which are highly ranked in terms of differential expression, but do not reach the threshold of FDR < 0.05. Therefore, to obtain a more complete view of the changes to gene expression in APOE4 or APOE2 mice relative to APOE3 mice, we performed enrichment analysis on the entire list of detectable genes using three methods of rank-based gene set enrichment analysis with different statistical methodologies: *fry* (Wu et al., 2010), *camera* (Wu and Smyth, 2012) and *GSEA*

(Subramanian et al., 2005) (implemented in the *fgsea* R package (Sergushichev, 2016)). We then combined raw p-values from each method to obtain an overall significance value by calculation of the harmonic mean p-value (Wilson, 2019) (a method which has been shown specifically to be robust to combining dependent p-values). After FDR adjustment of the harmonic mean p-value, we observed 73 gene sets to be significantly altered (FDR adjusted harmonic mean p-value < 0.05) in male APOE4 mice, 5 gene sets in male APOE2 mice, 30 gene sets in female APOE4 mice, and 30 gene sets in female APOE2 mice. This appears to be a considerable number of significantly altered gene sets, especially for young mice carrying mutations associated with LOAD. However, these effects are likely driven by both *APOE* genotype and litter-of-origin (**Figure 5**). Therefore, to simplify interpretation, we considered a gene set to be altered significantly if the FDR-adjusted harmonic mean p-value was < 0.01, which represents the most significantly altered pathways (as these are likely to have the largest effect of genotype). The statistical significance values of the gene sets are mostly driven by distinct expression signals, as shown by the lack of overlap of the DE genes found in the significantly altered gene sets (with the exception of the *KEGG* gene sets for oxidative phosphorylation and Parkinson's disease, which share 14 DE genes).

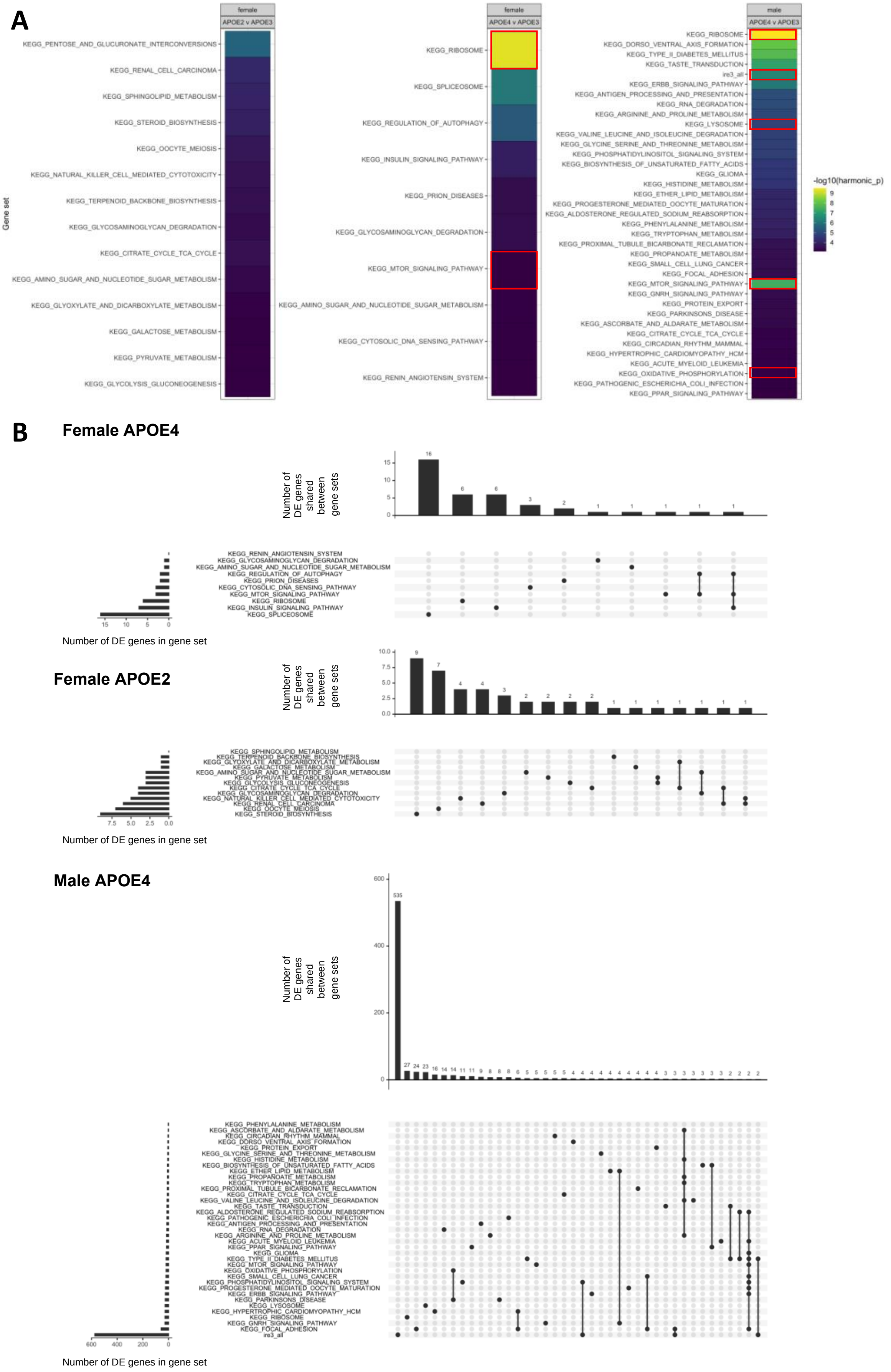
A
female
APOE2 v APOE3
female
APOE4 v APOE3
male
APOE4 v APOE3
Gene set
-log10(harmonic_p)
KEGG_PENTOSE_AND_GLUCURONATE_INTERCONVERSIONS
KEGG_RENAL_CELL_CARCINOMA
KEGG_SPHINGOLIPID_METABOLISM
KEGG_STEROID_BIOSYNTHESIS
KEGG_OOCYTE_MEIOSIS
KEGG_NATURAL_KILLER_CELL_MEDIATED_CYTOTOXICITY
KEGG_TERPENOID_BACKBONE_BIOSYNTHESIS
KEGG_GLYCOSAMINOGLYCAN_DEGRADATION
KEGG_CITRATE_CYCLE_TCA_CYCLE
KEGG_AMINO_SUGAR_AND_NUCLEOTIDE_SUGAR_METABOLISM
KEGG_GLYOXYLATE_AND_DICARBOXYLATE_METABOLISM
KEGG_GALACTOSE_METABOLISM
KEGG_PYRUVATE_METABOLISM
KEGG_GLYCOLYSIS_GLUCONEOGENESIS
KEGG_RIBOSOME
KEGG_SPLICEOSOME
KEGG_REGULATION_OF_AUTOPHAGY
KEGG_INSULIN_SIGNALING_PATHWAY
KEGG_PRION_DISEASES
KEGG_MTOR_SIGNALING_PATHWAY
KEGG_CYTOSOLIC_DNA_SENSING_PATHWAY
KEGG_RENIN_ANGIOTENSIN_SYSTEM
KEGG_DORSO_VENTRAL_AXIS_FORMATION
KEGG_TYPE_II_DIABETES_MELLITUS
KEGG_TASTE_TRANSDUCTION
ire3_all
KEGG_ERBB_SIGNALING_PATHWAY
KEGG_ANTIGEN_PROCESSING_AND_PRESENTATION
KEGG_RNA_DEGRADATION
KEGG_ARGININE_AND_PROLINE_METABOLISM
KEGG_LYSOSOME
KEGG_VALINE_LEUCINE_AND_ISOLEUCINE_DEGRADATION
KEGG_GLYCINE_SERINE_AND_THREONINE_METABOLISM
KEGG_PHOSPHATIDYLINOSITOL_SIGNALING_SYSTEM
KEGG_BIOSYNTHESIS_OF_UNSATURATED_FATTY_ACIDS
KEGG_GLIOMA
KEGG_HISTIDINE_METABOLISM
KEGG_ETHER_LIPID_METABOLISM
KEGG_PROGESTERONE_MEDIATED_OOCYTE_MATURATION
KEGG_ALDOSTERONE_REGULATED_SODIUM_REABSORPTION
KEGG_PHENYLALANINE_METABOLISM
KEGG_TRYPTOPHAN_METABOLISM
KEGG_PROXIMAL_TUBULE_BICARBONATE_RECLAMATION
KEGG_PROPANOATE_METABOLISM
KEGG_SMALL_CELL_LUNG_CANCER
KEGG_FOCAL_ADHESION
KEGG_GNRH_SIGNALING_PATHWAY
KEGG_PROTEIN_EXPORT
KEGG_PARKINSONS_DISEASE
KEGG_ASCORBATE_AND_ALDARATE_METABOLISM
KEGG_CIRCADIAN_RHYTHM_MAMMAL
KEGG_HYPERTROPHIC_CARDIOMYOPATHY_HCM
KEGG_ACUTE_MYELOID_LEUKEMIA
KEGG_OXIDATIVE_PHOSPHORYLATION
KEGG_PATHOGENIC_ESCHERICHIA_COLI_INFECTION
KEGG_PPAR_SIGNALING_PATHWAY
B
Female APOE4
Number of DE genes shared between gene sets
Number of DE genes in gene set
Female APOE2
Number of DE genes shared between gene sets
Number of DE genes in gene set
Male APOE4
Number of DE genes shared between gene sets
Number of DE genes in gene set

**Figure 8:** Ranked-list gene set enrichment testing. **A)** Heatmap indicating gene sets with a FDR-adjusted harmonic mean p-value < 0.01 in APOE-TR mice at 3 months of age. Gene sets of interest are highlighted with a red box. Note that no gene sets were found to contain an FDR-adjusted harmonic mean p-value of < 0.01 in male APOE2 mice. **B)** Upset plots indicating the overlap of DE genes across the gene sets which were calculated to have a FDR-adjusted harmonic mean p-value < 0.01 in APOE-TR mice.

As described in the main text, the *KEGG_OXIDATIVE_PHOSPHORYLATION* gene set was the only gene set to be affected by all the EOfAD-like mutations in our zebrafish models and not by non AD-relevant mutations. In APOE-TR mice, statistical evidence was observed for this gene set to be altered in male APOE4 mice (**Table 1**). The overall direction of logFC of genes in the *KEGG_OXIDATIVE_PHOSPHORYLATION* gene set was up in the male APOE4 mice (although some genes of the gene set were downregulated) (**Figure 9**).

**Table 1: Significance of the *KEGG_OXIDATIVE_PHOSPHORYLATION* gene set in young APOE-TR mice.**

| Sex | APOE | FDR-adjusted harmonic mean p-value |
|---|---|---|
| Male | APOE4 | 0.00948 |
| | APOE2 | 0.794 |
| Female | APOE4 | 0.794 |
| | APOE2 | 0.248 |

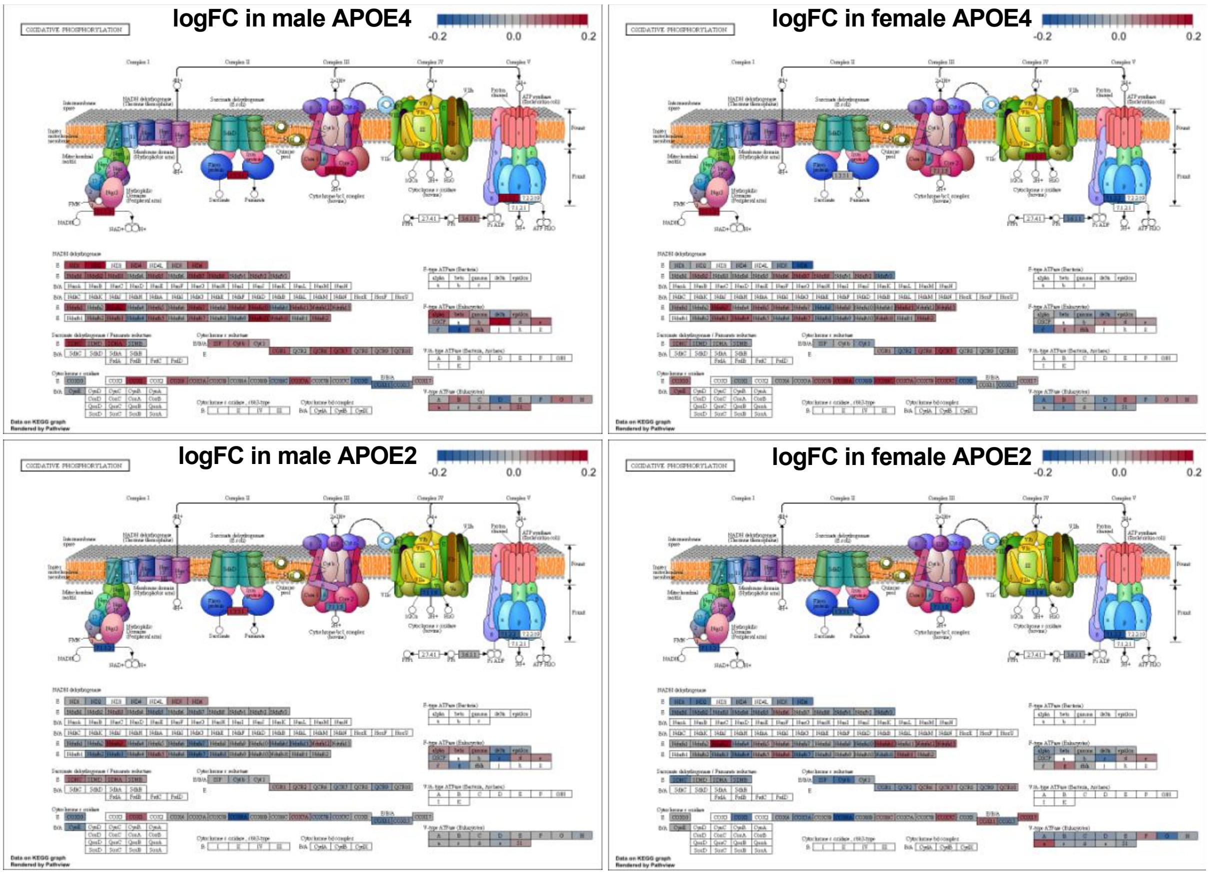


**Figure 9:** Changes to the *KEGG_OXIDATIVE_PHOSPHORYLATION* gene set in young APOE-TR mice.

Additionally, the *KEGG_RIBOSOME* gene set was found to be altered in common by EOfAD-like (and non-EOfAD-like) mutations in our zebrafish models. This gene set was found to be highly significantly altered in both male and female APOE4 mice and not in APOE2 mice (**Figure 8A**). The genes in this gene set appear to be mostly upregulated in APOE4 mice, and are both upregulated or downregulated in APOE2 mice.

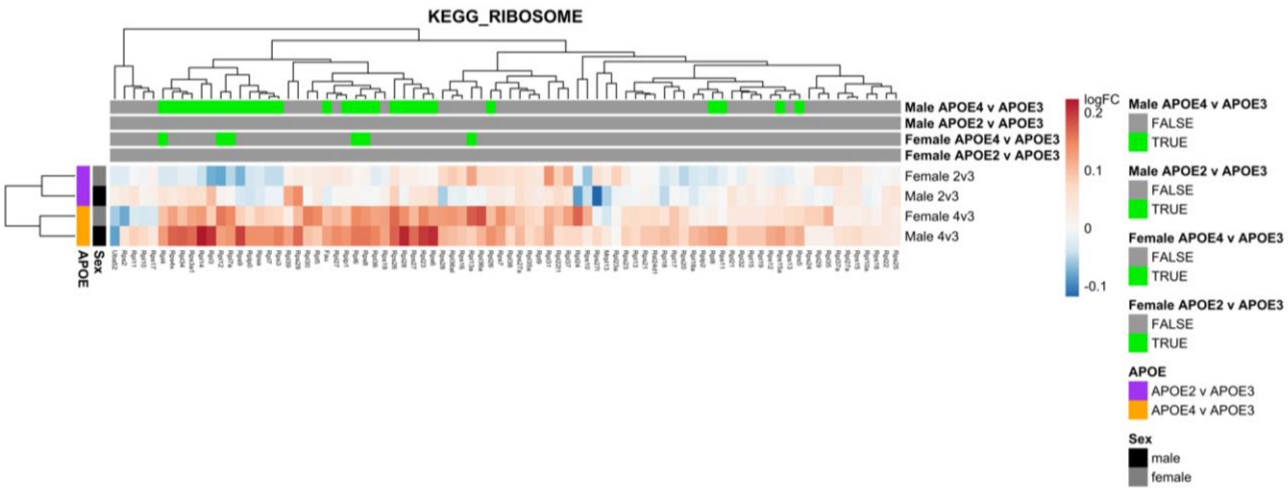


**Figure 10:** The columns in the heatmap represent a gene in the *KEGG_RIBOSOME* gene set, while the rows indicate the comparisons between APOE-TR mice in the differential expression analysis. The colour of s cell represents the logFC of a particular gene, and the genes are labelled in green above if they were classified as differentially expressed (FDR < 0.05) in the differential gene expression analysis in a particular comparison.

The *KEGG_MTOR_SIGNALING_PATHWAY* gene set was observed to be significantly altered in both male and female APOE4 mice. No clear direction of change is evident, as both up- and down-regulation of genes in this pathway is observed (**Figure 11**).

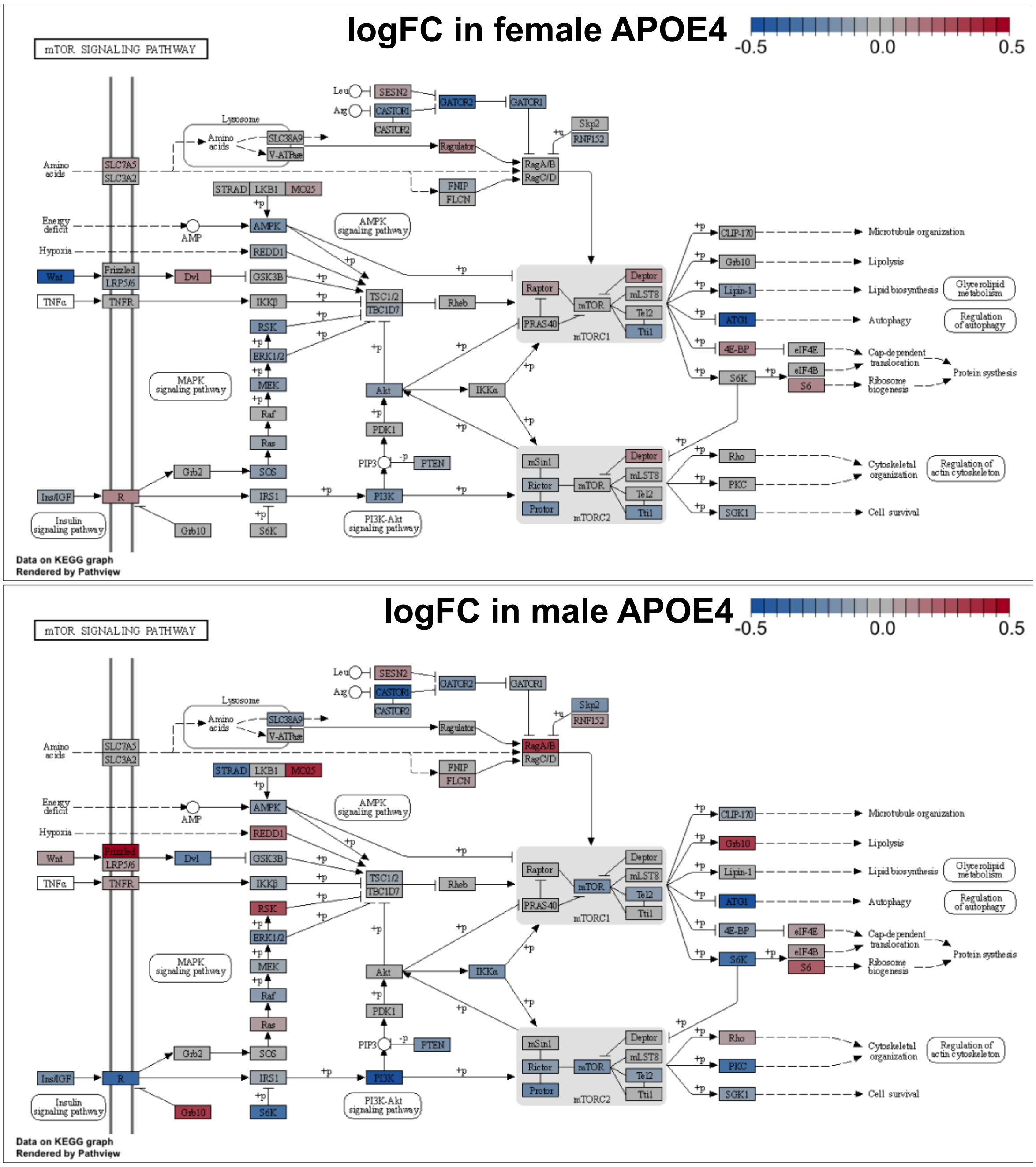


Figure 11: Changes to the *KEGG_MTOR_SIGNALING_PATHWAY* gene set in young APOE4 mice.

### Cell type proportions

We next assessed whether changes to gene expression observed in young APOE-TR mice are due to changes to cell type proportions as described in **Supplementary File 2**. We observed a slight decrease in expression values of neuronal marker genes in female APOE2 mice, and increased expression of marker genes of oligodendrocytes and astrocytes in male APOE4 mice, suggesting that the overall changes to gene expression observed in these mice may be driven partially by differences in cell type proportions (**Figure 12**).

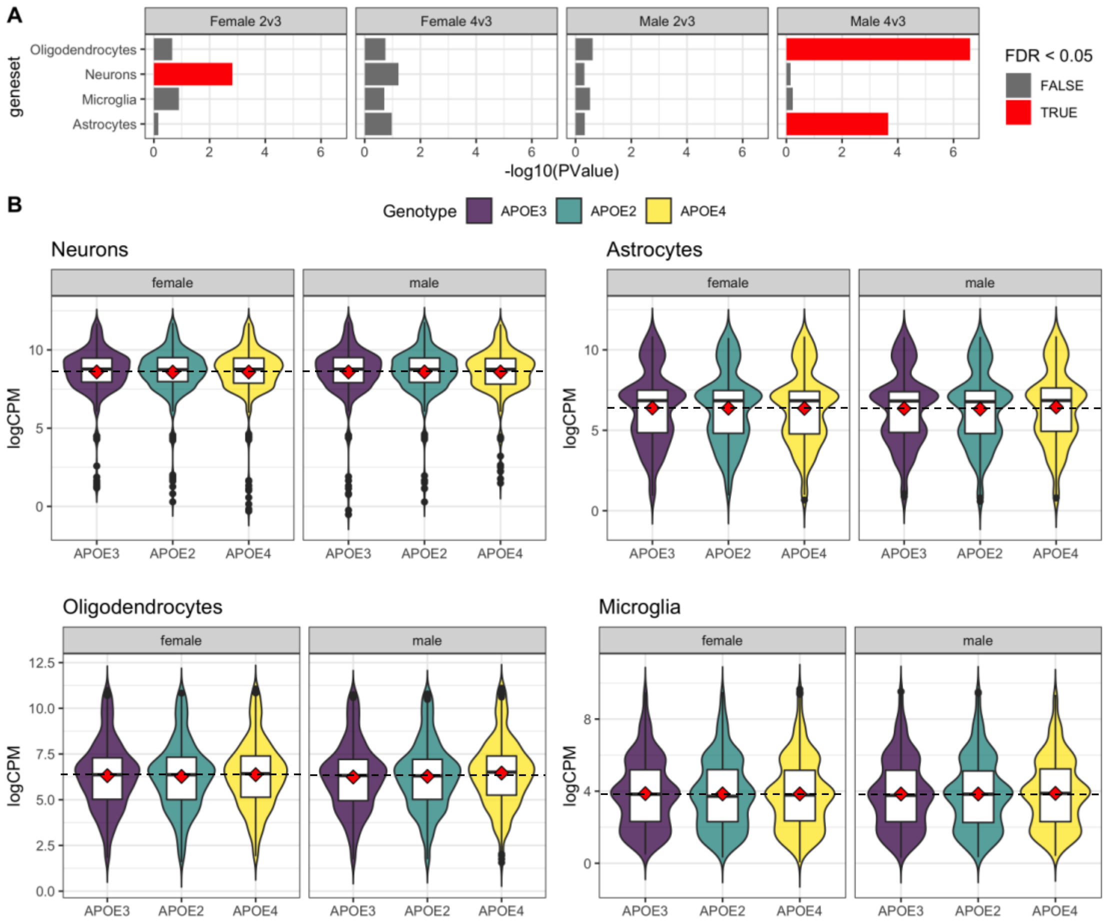


**Figure 12: A)** Significance of gene set testing from *fry* with a directional hypothesis of gene sets consisting of marker genes of neurons, astrocytes and oligodendrocytes from (Cahoy et al., 2008) and microglia from (Oosterhof et al., 2017). **B)** Distribution of expression (logCPM) of these cell type marker genes in APOE-TR mice. Boxplots show the summary statistics, while the violin plots summarise the density of logCPM expression values. The mean of each experimental group is shown by the red diamonds, and the mean of the APOE3 expression values is shown by black dashed lines.

### Discussion and conclusion

In conclusion, our re-analysis of the comprehensive study of Zhao et al. (2020) predicts many cellular processes to be affected in young APOE-TR mice. However, sources of external variation may be contributing to the observed effects (i.e. litter-of-origin, changes to cell type proportions and possibly gene length). Importantly, the specific changes to gene expression observed in our zebrafish models of EOfAD are observed to be altered in APOE4 mice (*KEGG_RIBOSOME* in both males and females, and *KEGG_OXIDATIVE_PHOSPHORYLATION* only in males). However, replication of this study is desirable, particularly with modifications to compensate for the litter-of-origin confounding effect, to confirm that these effects are due to *APOE* genotype.
