## Supplemental File 4 for "Comparative analysis of Alzheimer’s disease knock-in model brain transcriptomes implies changes to energy metabolism as a causative pathogenic stress"

**Energy production changes in zebrafish and mouse models of genetic variation driving AD**

The majority of heterozygous EOfAD-like mutations we have studied in zebrafish show overall (majority) downregulation of the oxidative phosphorylation gene set in young adult brains relative to brains of the wild type genotype. Only heterozygosity for the T141_L142delinsMISLISV (reading frame-preserving) mutation of *psen2* has been seen to give overall upregulation of these genes. Another complex, yet probably EOfAD-like mutation in *psen2* we have studied in zebrafish: *psen2*$^{S4ter}$, (which likely produces Psen2 proteins lacking N-terminal sequences) also showed strong overall upregulation of the oxidative phosphorylation gene set (Jiang et al., 2020), although that dataset contains technical artefacts which complicate interpretation and so it was not included in this paper. Transheterozygosity for mutations in *sorl1* also results in overall upregulation of oxidative phosphorylation genes. We are uncertain as to why this variability in effects on the oxidative phosphorylation gene set occurs. It may be that the disruption of this gene set that is consistently observed is a product of both genotype and environmental factors, i.e. the mutant fish may be more or less responsive to environmental variation such changes in water quality, microbiome, handling etc.. Also, it can be misleading to infer the direction of change in a particular cell activity, such as oxidative phosphorylation, based on the majority behaviour of a (somewhat arbitrarily) defined set of genes. Obviously, actual measurement of e.g. respiratory rates in the zebrafish mutant brains would be needed to establish, with certainty, how the mutations are affecting oxidative phosphorylation. Note, however, that the subtlety of the gene regulatory effects we have observed in the fish models means that discernment of physiological oxygen consumption differences between mutant fish and their wild type siblings may be challenging. (Simultaneous measurement of differences in the expression levels of the approximately 100 genes in the oxidative phosphorylation gene set gives great sensitivity for detection of statistically significant differences.)

Importantly, male mice homozygous for the late onset AD (LOAD) risk allele, APOE4, showed altered expression of the oxidative phosphorylation gene set, while female APOE4 mice showed a similar trend that did not reach the threshold for statistical significance. This demonstrates the similarity, at the molecular level, between the cellular effects of genetic variants causing EOfAD and promoting

LOAD, and supports the validity of analysing early molecular events in AD pathogenesis using knock-in models in zebrafish. By extension, our results also support that ‘omics analyses of the brains of mouse knock-in models of EOfAD mutations will yield valuable information on AD pathogenesis. Although numerous such models were created 15-20 years ago, and some showed subtle cognitive effects (Guo et al., 1999; Kawasumi et al., 2004), their relevance to understanding AD was apparently dismissed since they failed to reproduce the Aβ plaque and neurofibrillary tangle phenotypes of the human disease. Consequently, the molecular state of their brains has never been analysed in detail.

In humans, it is thought that Aβ-related abnormalities precede metabolic defects (Jack et al., 2013; Leclerc and Abulrob, 2013). However, metabolic abnormalities are generally measured *in vivo* using FDG-PET imaging, which is likely not sensitive enough to detect subtle changes. Nevertheless, reduced glucose uptake has been observed by FDG-PET in the brains of living subjects before the onset of dementia (Mosconi et al., 2008; Mosconi et al., 2009). Additionally, changes to the expression of genes in the oxidative phosphorylation pathway has been observed in the post-mortem brains of early, and late AD subjects relative to age-matched controls (Manczak et al., 2004). Neuronal cells derived from human induced pluripotent stem cells (hiPSCs) of LOAD patients also show increased expression of oxidative phosphorylation proteins and oxidative stress (Birnbaum et al., 2018). While neurons derived from a patient carrying the $\textit{PSEN1}^{S170F}$ EOfAD mutation also show mitochondrial abnormalities (Li et al., 2020). Together, these results support that changes to mitochondrial function are an early AD pathology.

Our findings regarding EOfAD mutations in zebrafish were not consistent with findings from our analysis of transcriptome data from  in homozygous $\textit{App}^{NL\text{-}G\text{-}F}$ mice. However, this transcriptome data was generated from “middle-aged” (12 month old) mice rather than young adults, and the endogenous *App* gene of the mouse was altered with a total of six mutations (three that humanise the sequence for the Aβ region and three EOfAD mutations), motivated by the idea that the more aggregation-prone human Aβ sequence plays a critical role in the pathogenic mechanism of AD. Therefore, it is not directly comparable with our zebrafish EOfAD models, which contain single, EOfAD-like mutations within single alleles of endogenous genes. To our knowledge, a transcriptome analysis has not been performed on this model at a younger age. However, the expression of genes involved in lysosomal function (*KEGG_LYSOSOME*) was observed to be highly significantly upregulated in the homozygous *App*$^{NL\text{-}G\text{-}F}$ brains. This is not unexpected, as acidification of the endo-lysosomal system is impaired by increased levels of the β-CTF fragment of APP (also known as C99 and generated by β-secretase cleavage product of APP (Jiang et al., 2019)). Increased β-CTF has been observed in the brains of *App*$^{NL\text{-}G\text{-}F}$ mice (Saito et al., 2014). In a mouse model of a lysosomal storage disorder (Glycogen storage disease type 2, which most seriously affects muscle (Yambire et al., 2019)) lysosomes failed to become sufficiently acidic and this resulted in an intracellular ferrous iron deficiency and a pseudo-hypoxic response (degradation of HIF1-α, the master transcriptional regulator of the cellular response to hypoxia, is dependent on both oxygen and ferrous iron (Ivan et al., 2001)), mitochondrial dysfunction and inflammation (Yambire et al., 2019). Additionally, the *App*$^{NL\text{-}G\text{-}F}$ mouse model shows increased levels of Aβ from a young age (Saito et al., 2014), and the deposition of Aβ into plaques was shown to be associated with the increased expression of genes in the complement system in the comprehensive spatial transcriptomics study with aging in the *App*$^{NL\text{-}G\text{-}F}$ mouse model (Chen et al., 2020), providing another avenue for these mutations to trigger inflammation. Mitochondrial dysfunction has not been observed directly in *App*$^{NL\text{-}G\text{-}F}$ mice. However, increased levels of oxidative stress have been observed at 12 months of age (Izumi et al., 2020), suggestive of increased reactive oxygen species (ROS) that can be generated by dysfunction of mitochondrial respiration. Therefore, we suspect that similar processes are being affected in *App*$^{NL\text{-}G\text{-}F}$ mice to those in our zebrafish models and in APOE4 mice. However, their subtle signs in the transcriptome may be obscured by noise from the strong inflammatory signals in the bulk brain transcriptomic data (as well as confounding influences on the transcriptome analysis such as sex and litter-of-origin effects).

**mTOR signalling can regulate ribosomal gene set expression**

In the majority of our zebrafish mutants (all except for zebrafish transheterozygous for EOfAD-like mutations in *sorl1*) and in the APOE4 mouse data we have observed changes to the expression of the set of genes encoding components of the ribosomal subunits. Protein translation is one of the most energy-costly processes within a cell (Buttgereit and Brand, 1995), and so expression of ribosomal proteins is modulated by the mammalian target of rapamycin (mTOR) system that surveys cellular nutrient status to adjust cellular metabolism (reviewed in (Mayer and Grummt, 2006; Zhou et al., 2015)). mTOR signalling, which appears to be increased in late-stage AD brains compared to controls (Griffin et al., 2005; Li et al., 2005; Sun et al., 2014), is regulated by growth factors, nutrients, energy levels and stress. In addition to ribosome biogenesis, mTOR signalling plays a role in various cellular processes which are also implicated in AD pathogenesis, such as autophagy and metabolism (reviewed in (Saxton and Sabatini, 2017)).

The mTOR proteins reside at lysosomes within the mTORC1 and mTORC2 protein complexes (Sancak et al., 2010). Intriguingly, the v-ATPase complex that acidifies the endolysosomal pathway is required for mTORC1 activation (Zoncu et al., 2011). Proper assembly of the v-ATPase at the lysosome requires the PSEN1 protein (and this process is impaired in EOfAD patient fibroblasts) (Lee et al., 2010). Stimulation of mTOR signalling has been observed in response to accumulation of Aβ (Caccamo et al., 2010), while hyperactivation of mTOR is observed in Down's syndrome (where the dosage of the *APP* gene is increased because it resides on Chromosome 21 and early onset AD is common) (Bordi et al., 2019; Iyer et al., 2014). Intriguingly, Bordi et al. (2019) observed that inhibition of mTOR signalling (specifically, mTORC1) restores auto- and mito-phagy defects in fibroblasts from Down's syndrome individuals. Among our transcriptome analyses of AD models, we only observed statistically significant changes to the expression of genes in the *KEGG* gene set for mTOR signalling in transheterozygous *sorl1* mutants, in *psen1*$^{Q96\_K97del}$/+ mutant zebrafish after acute hypoxia exposure, and in both male and female APOE4 mice. However, the majority of regulation of mTOR signalling occurs at the protein level, so that it is perhaps unsurprising that we could only detect significant changes in the transcriptional response to altered mTOR signalling in the other mutants.

**Advantages and disadvantages of zebrafish for analysis of genetic variants driving AD**

In a highly sensitive analysis method such as RNA-seq, external sources of variation must be minimised. Our analysis has revealed that zebrafish can be highly advantageous for transcriptome profiling in the context of RNA-seq, as large numbers of progeny can be produced from a single pair mating, and these can subsequently be raised together in a single aquarium system, thus reducing both genetic and environmental sources of variation. This has allowed us to observe subtle effects due to the EOfAD-like mutations we have analysed. In contrast, a female mouse can only birth small litters of 5-10 pups, making it particularly difficult to obtain sufficient samples which are synchronous siblings (particularly when a genotype of interest is homozygous). Our re-analysis of the APOE-TR mouse brain transcriptomes was unable to distinguish with great certainty whether the effects we observed were due to *Apoe* genotype or litter-of-origin. Also, information of whether $App^{NL\text{-}G\text{-}F}$ mice were littermates was not available, and this required us to assume that the effect of litter was negligible in order to perform the analysis. Another contrast between brain transcriptome analysis in zebrafish compared to mice is the influence of sex. Mouse brain transcriptomes show very significant difference due to sex, while this has a negligible effect on bulk brain transcriptomes from zebrafish (Barthelson et al., 2020a; Barthelson et al., 2020b; Barthelson et al., 2020c), and can generally can be ignored in a differential expression analysis. We also found evidence for changes to cell type proportions in both APOE-TR and $App^{NL\text{-}G\text{-}F}$ mice, a phenomenon that can create the artefactual appearance of gene expression change. We have not observed cell-type proportion differences in young or “middle aged” (24 month old) zebrafish (Barthelson et al., 2021; Barthelson et al., 2020a; Barthelson et al., 2020b; Barthelson et al., 2020c; Hin et al., 2020), possibly associated with the resistance to damage of the highly regenerative zebrafish brain (Kroehne et al., 2011). While this regenerative ability may hinder use of zebrafish for studying overt neurodegeneration, it can facilitate analysis of young, bulk brain transcriptomes before overt pathological processes would be expected.

The advantages of zebrafish for analysing the early effects of EOfAD mutations are countered, occasionally, by disadvantages. The teleost lineage in which zebrafish arose from underwent a whole-genome duplication event (reviewed in (Meyer and Van de Peer, 2005)), such that many human genes are represented by duplicate “co-orthologues” in zebrafish (e.g. the co-orthologues of *APP* and *APOE* in zebrafish are *appa / appb* and *apoea / apoeb* respectively). This complicates interpretation of the effects of mutations in these genes. Additionally, zebrafish have never been shown, definitively, to be capable of producing Aβ, a pathological hallmark of AD. The β-secretase (BACE) site of human APP does not appear conserved in zebrafish Appa and Appb (Moore et al., 2014). Whether Aβ is causative of AD pathology or a consequence of it continues as a matter of debate in the AD research community (reviewed in (Morris et al., 2018)). If zebrafish cannot produce Aβ, then the changes we have observed in the brains of our zebrafish models may illuminate Aβ-independent effects of EOfAD mutations.
